# Heterogeneity of satellite cells implicates DELTA1/NOTCH2 signaling in self-renewal

**DOI:** 10.1101/824359

**Authors:** Valeria Yartseva, Leonard D. Goldstein, Julia Rodman, Lance Kates, Mark Z. Chen, Ying-Jiun J. Chen, Oded Foreman, Christopher W. Siebel, Zora Modrusan, Andrew S. Peterson, Ana Jovičić

## Abstract

How satellite cells and their progenitors balance differentiation and self-renewal to achieve sustainable tissue regeneration is not well understood. A major roadblock to understanding satellite cell fate decisions has been the difficulty to study this process *in vivo*. By visualizing expression dynamics of myogenic transcription factors during early regeneration *in vivo*, we identified the time point at which cells undergo decisions to differentiate or self-renew. Single-cell RNA sequencing revealed heterogeneity of satellite cells during both muscle homeostasis and regeneration, including a subpopulation enriched in *Notch2* receptor expression. Furthermore, we reveal that differentiating cells express the *Dll1* ligand. Using antagonistic antibodies we demonstrate that the DLL1 and NOTCH2 signaling pair is required for satellite cell self-renewal. Thus, differentiating cells provide the self-renewing signal during regeneration, enabling proportional regeneration in response to injury while maintaining the satellite cell pool. These findings have implications for therapeutic control of muscle regeneration.

## INTRODUCTION

Skeletal muscle possesses remarkable regenerative capacity mediated by resident stem cells, called satellite cells (Hawke & Garry 2001; Mauro 1961). Throughout life satellite cells are tasked with muscle fiber maintenance and repair through proliferation, differentiation and self-renewal. Maintaining an appropriate balance between differentiation and self-renewal is critical for preserving the stem cell pool over time and for proportional tissue regeneration upon injury. While tremendous efforts have elucidated mechanisms of muscle differentiation (Braun & Gautel 2011), signals that mediate self-renewal have remained elusive (Zammit 2008; Giordani et al. 2018).

Satellite cells, defined by expression of the Paired Box 7 (PAX7) transcription factor, in homeostatic muscle reside in a quiescent state between the basal lamina and the muscle fiber (Mauro 1961; Seale et al. 2000). Upon muscle injury or disease, satellite cells become activated to enter the cell cycle, upregulate Myogenic factor 5 (MYF5) (Beauchamp et al. 2000) and Myoblast determination protein (MYOD1) (Cornelison & Wold 1997), and proliferate to generate myoblast progenitors (Braun & Gautel 2011; Zammit et al. 2006). As activated cells progress along the differentiation path, they begin expressing Myogenin (MYOG) (Hasty et al. 1993), an indication of differentiation commitment. Importantly, concomitant with differentiation, satellite cells need to self-renew to replenish the stem cell pool (Collins et al. 2005; Zammit et al. 2004). One of the fundamental questions in regenerative biology that remains unanswered is how do satellite cells balance proliferation, differentiation and self-renewal?

The ability to investigate the mechanisms regulating self-renewal depends on the ability to study cells as they undergo cell fate decisions. These functions have typically been interrogated in myogenic cells propagated on artificial 2D substrates *in vitro* or single myofiber explants *ex vivo* (Fujimaki et al. 2018; Kuang et al. 2007; Zammit 2008). However, myogenic cells under these conditions do not fully recapitulate innate satellite cell behavior, nor the complex environment of muscle tissue and their utility as models of satellite cell fate decision and muscle regeneration is limited (Webster et al. 2016). For these reasons, investigating cells in their native environment as they undergo cell fate decisions is the first step in elucidating specific physiological signals that directs self-renewal.

Here we developed a novel *in vivo* paradigm to study satellite cell fate decisions. We utilized RNAscope *in situ* hybridization technology to visualize both the timing and spatial expression patterns of *Pax7* (satellite cell marker), *Myod1* (activation and proliferation marker) and *Myog* (early differentiation marker) during early phases of cardiotoxin-induced regeneration. By doing so, we identified the time point at which satellite cells undergo cell fate decisions *in vivo* and were able to capture cells as they became committed to differentiation or maintained their self-renewal capacity. Single-cell RNA sequencing revealed heterogeneity among myogenic cells, reflecting functional roles and dynamic states required for successful regeneration. We show that along the differentiation path, myogenic cells downregulate Notch receptors, while upregulating the Notch ligand *Delta like canonical Notch ligand 1* (*Dll1*). By employing selective antagonist antibodies to inhibit individual Notch receptors or ligands during regeneration *in vivo*, we demonstrate that inhibition of DLL1 activity results in the loss of *Pax7*-positive progeny and impairs satellite cell self-renewal. Furthermore, we identify NOTCH2 as the principal regulator mediating self-renewal. Our data suggest a model whereby differentiating DLL1-expressing cells convey a self-renewing signal to satellite cells, thus ensuring proportional muscle regeneration.

## RESULTS

### Myogenic transcription factor dynamics during muscle injury

To better understand the dynamic changes in satellite cells (SCs) and descendant myogenic progenitors upon injury *in vivo*, we performed a time course analysis of myogenic transcription factor expression during early stages of cardiotoxin-induced regeneration. We used RNAscope *in situ* hybridization for direct visualization of expression dynamics of *Pax7*, *Myod1* and *Myog* from 0 to 4 days post-injury in mouse muscle tissue sections (Figure 1 and S1). We simultaneously labeled *Pax7* and *Myod1*, or *Pax7* and *Myog*. Labeling with *Pax7* and *Myod1* probes revealed that activated satellite cells co-express these two markers at earliest stages of regeneration (days 1–2.5). *Pax7*+/*Myod1*– cells were rarely observed. During this stage, *Myog* was rarely detected. At day 3 post-injury we uncovered notable changes in expression patterns. *Myod1* and *Pax7* signals started being largely separated, indicating that these mRNAs were now enriched by different cells (Figure S1). This onset of *Pax7* and *Myod1* separation among cells coincided with a strong increase in *Myog* expression. Such pattern of *Pax7*, *Myod1* and *Myog* expression continued into day 4. This indicated that at day 3 post-injury myogenic cells reached cell fate decision stage, yielding cells that were either committed to differentiation or maintenance of the capacity to self-renew. Thus day 4 post-injury offers an ideal time point to interrogate the mechanisms regulating satellite cell fate decisions and the two different states.

**Figure 1.**
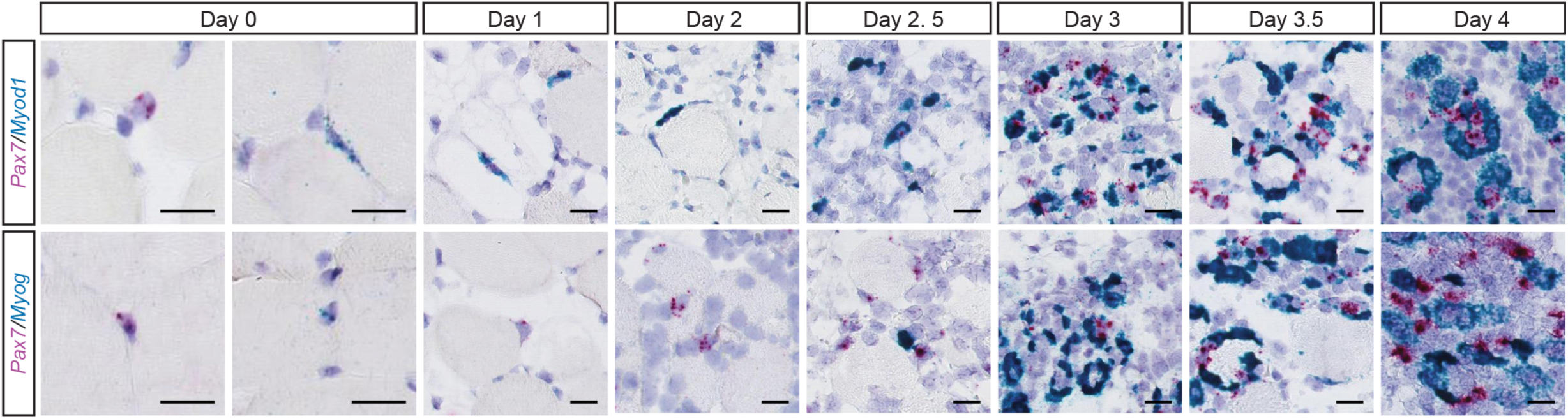
*In situ* analysis of myogenic transcription factor expression in early muscle regeneration. Representative images of *in situ* hybridization showing expression of myogenic factors *Pax7*, *Myod1* and *Myog* in cardiotoxin-injured tibialis anterior (TA) muscle at indicated time points. *Pax7* probe was visualized in pink and was used on every section, both with *Myod1* or *Myog* probes (blue). Strong *Myod1* signal co-localized with *Pax7* signal at days 1, 2, 2.5, and *Pax7+*/*Myod1*-cells were rarely detected. In serial sections, co-labeling for *Pax7* and *Myog*, revealed that *Myog*+ cells are rarely detected at days 1, 2, 2.5, instead singly *Pax7+* cells are readily detected. Starting at day 3 a notable upregulation of *Myog* was observed. Scale bar is 20 μm.

### Single-cell RNA sequencing reveals satellite cell heterogeneity

To gain insight into the potential mechanisms regulating the observed cellular states and cell fate decisions during muscle regeneration, we performed single-cell RNA sequencing of satellite cells during homeostasis (day 0) and at day 4 post-cardiotoxin injury.

First, we analyzed expression profiles for 2,062 satellite cells isolated from hind limbs during muscle homeostasis (day 0). To characterize different cellular states during homeostasis, we performed unsupervised clustering, which separated cells into two clusters (Figure 2A and B, Table S1). The first cluster showed no strong enrichment for individual genes, but was depleted of *Myf5* expression (Figure S2A), suggesting these cells are quiescent satellite cells with a basal expression program or low transcriptional activity. By contrast, the second cluster was characterized by increased expression of specific genes, including higher expression of *Sdc4*, previously associated with higher self-renewing capacity (Ono et al. 2012), and *Notch2*, a Notch receptor (Figure 2B). To validate the existence of the *Sdc4/Notch2*-enriched subpopulation of satellite cells in homeostasis, we analyzed an independently generated and previously published dataset (Giordani et al. 2019), which profiled skeletal muscle-resident cell populations during homeostasis. By applying the same workflow for satellite cell selection and clustering as with our dataset, we detected a *Sdc4/Notch2-*enriched subpopulation (Figure S2B and C) and confirmed a high overlap of cluster marker genes between the two independently generated datasets (Figure S2D).

**Figure 2.**
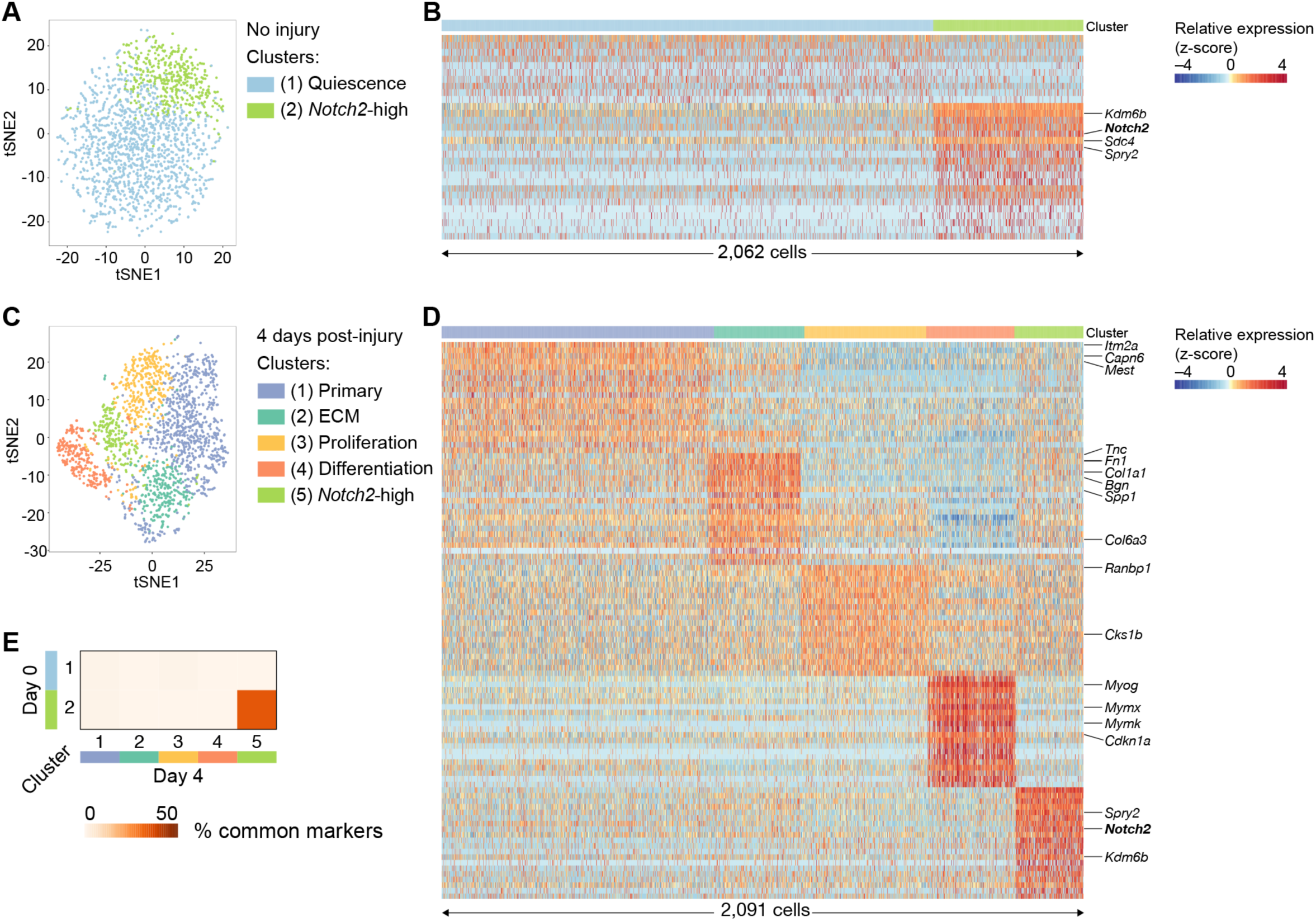
Single-cell RNA sequencing reveals satellite cell heterogeneity. (A) t-distributed stochastic neighbor embedding (t-SNE) of single-cell expression profiles at day 0 (n = 2,062). Cells clustered into two clusters: 1 ‘Quiescence’ (n = 1,580), 2 ‘*Notch2*-high’ (n = 482). (B) Heatmap of top identified marker genes for clusters shown in (A). Shown are all 10 identified markers for cluster 1 and the top 20 markers for cluster 2. Columns and rows correspond to cells and genes, respectively. Cells are ordered by cluster as shown in top sidebar, within-cluster order is random. (C) t-SNE plot of single-cell expression profiles at day 4 (n = 2,091). Cells clustered into five clusters: 1 ‘Primary’ (n = 868), 2 ‘ECM’ (n = 302), 3 ‘Proliferation’ (n = 414), 4 ‘Differentiation’ (n = 287), 5 ‘*Notch2*-high’ (n = 220). (D) Heatmap of top 20 identified marker genes for clusters shown in (C), otherwise as in (B). (E) Cluster comparison based on common markers (Jaccard index).

Next, we analyzed expression profiles for 2,091 myogenic cells isolated from tibialis anterior (TA) at 4 days post cardiotoxin-injury (day 4). Unsupervised clustering identified five clusters consistent with active roles in self-renewal, proliferation and differentiation (Figure 2C, D, Table S2). Part of the observed heterogeneity among myogenic cells could be explained by different stages of differentiation. Clusters 1, 2 and 5 were enriched in *Pax7* expression, while clusters 3 and 4 were enriched in *Myod1* expression (Figure S2E). In addition to *Myod1*, cells in cluster 3 were enriched for cyclin genes (*Cks1b, Ccna2, Cdk4, Ccnb2*), mitochondrial components and oxidative phosphorylation, thus likely representing transit-amplifying myoblast cells (‘proliferation’ cluster). Cluster 4 cells were characterized by muscle differentiation gene expression such as *Myog*, *Mymx*, *Mymk*, *Hes6* (Cornelison & Wold 1997; Hasty et al. 1993; Bi et al. 2018; Gao et al. 2001; Cossins et al. 2002), and genes that act to suppress the cell cycle and mitosis (*Cdkn1a* and *Cdkn1c*), indicating that these cells are committed to differentiation (‘differentiation’ cluster). Interestingly, the differentiation cluster was also characterized by expression of *Delta like canonical Notch ligand 1* (*Dll1*) expression and diminished expression of Notch receptor genes, indicating that there is a ligand-biased myogenic cell lineage commitment.

Clusters 1, 2 and 5 were characterized by enrichment in *Pax7* expression and diminished *Myod1* compared to clusters 3 and 4. These three clusters revealed heterogeneity within the satellite cell compartment and pointed to different roles during regeneration. Cluster 1 showed higher expression of *Pax7* compared to clusters 2 and 5. During homeostasis satellite cells expressing higher levels of *Pax7* are less primed for myogenic commitment compared to cells with low *Pax7* levels (Rocheteau et al. 2012). This suggests that cluster 1 cells are most closely related to quiescence and retain their capacity to self-renew during regeneration (‘primary’ cluster). Among other genes enriched in cluster 1 were *Itm2a*, a known PAX3 target gene expressed during muscle development and in satellite cells (Lagha et al. 2013; Der Vartanian et al. 2019; Scaramozza et al. 2019), *Capn6* suppressor of myogenic differentiation (Tonami et al. 2013) and *Mest* implicated in muscle growth and regeneration (Hiramuki et al. 2015). Cluster 2 was enriched for numerous genes encoding extracellular matrix proteins, such as *Fn1*, collagen family I, III, V and VI members, *Tnc*, *Bgn* and *Spp1* (‘ECM’ cluster). Collagen type V and fibronectin 1 have been shown to regulate satellite cells by creating the appropriate niche to maintain their quiescence (Baghdadi et al. 2018; Bentzinger et al. 2013). This suggests that this cell subpopulation regulates self-renewal through ECM synthesis. Cluster 5 was specifically enriched in *Notch2* expression (‘*Notch2*-high’ cluster), with cluster markers similar to the ‘*Notch2*-high’ cluster from day 0 (Figure 2E) and the *Notch2-*enriched cluster identified in the (Giordani et al. 2019) dataset (Figure S2F). Additionally, *Notch2*-high cells appeared to be enriched for the G1 phase of the cell cycle, marked by increased expression of *Cdkn1b* and *Cdk6* (Figure S2E). Interestingly, this cluster showed reduced expression of *Notch3*, with *Notch2* and *Notch3* showing inverse expression patterns across the primary, ECM and *Notch2-*high clusters.

In summary, we observed two subpopulations of satellite cells during homeostasis and five subpopulations of myogenic cells during regeneration, including three subpopulations of activated satellite cells. Intriguingly, we uncovered variability in the expression of Notch components, with a subpopulation characterized by high *Notch2* expression that persists during homeostasis and regeneration, and a differentiating subpopulation characterized by low expression of Notch receptors and high expression of the *Dll1* ligand.

### Subpopulations represent dynamic satellite cell states

The identified clusters did not show strong separation (Figure 2C) and cells in one cluster often expressed markers for another cluster, albeit at lower levels (Figure S3A). These observations are consistent with clusters representing dynamic states of myogenic cells. For example, individual cells in the *Notch2*-high cluster showed moderate levels of expression for gene sets that typify one (and only one) of the other four cell populations (Figure S3A). Such expression patterns could be attributable to cells of one category transitioning through or into one of the other cell states. To account for intermediate expression profiles due to transitions between cellular states, we complemented the clustering analysis with a pseudotime analysis, which orders cells on a continuous trajectory of gene expression changes (Trapnell et al. 2014). Pseudotime analysis at day 4 post-injury constructed a trajectory with three branches (Figure S3B–F). Proliferating and *Notch2*-high cells were enriched on branch 1, primary and ECM cells on branch 2, and differentiating cells on branch 3. Co-localization of *Notch2*-high and proliferating cells, together with the observation that *Notch2-*high cells are enriched for the G1 phase of the cell cycle, suggest that *Notch2*-high cells may represent a transition state with a role in deciding cell fate. In addition, co-localization of primary and ECM cells supports the previously described role of extracellular matrix components synthesized by satellite cells in regulating quiescence.

### DLL1- and NOTCH2-enriched subpopulations exist *in vivo*

The intriguing observation of heterogeneity among satellite cells during homeostasis and regeneration, prompted us to further corroborate the observed subpopulations and confirm that they represent biological variation that is seen *in vivo*.

First, we co-labeled non-injured muscle tissue sections with a PAX7 antibody to mark satellite cells and an *in situ* hybridization probe recognizing *Notch2* mRNA. We found that within PAX7-positive cells *Notch2* expression was high in ∼20%, low in ∼50%, and absent in ∼30% of satellite cells (Figure S4A), which validates the heterogeneity in *Notch2* expression during homeostasis uncovered by single-cell RNA sequencing.

Second, we sought to confirm heterogeneity among satellite cells at day 4, as well as the upregulation of Notch ligand *Dll1* upon commitment to differentiation. We performed *in situ* hybridization analyses of tissue expression of *Itm2a* (primary cluster) and *Sdc4* (*Notch2*-high cluster). Only a subset of *Pax7+* satellite cells showed enrichment in each of these markers (Figure S4B and C). Importantly, when co-labeled *Itm2a* and *Sdc4* localized to different cells (Figure S4D). In addition, immunostaining for NOTCH2 on *PAX7.Cre_tdTomato* tissue sections detected both NOTCH2+ and NOTCH2– tdTomato cells (Figure S4E). Interestingly, among the NOTCH2+ cells we detected both cells with cytoplasmic or nuclear signal, indicating active NOTCH2-signaling. To validate the heterogeneity of ECM marker expression, we performed immunohistochemistry (IHC) on *PAX7.Cre_tdTomato* sections to visualize SPP1 and detected both SPP1+ and SPP1-tdTomato+ cells (Figure S4F).

Our single-cell RNA-sequencing results showed that commitment to differentiation is accompanied by a switch from Notch receptor expression to Dll1 ligand expression. ISH analysis confirmed that *Dll1* was co-expressed with *Myog* and not *Pax7* (Figure S4G, H). Importantly, genes enriched in non-differentiating subpopulations such as *Itm2a* or *Notch2* were not co-expressed with *Myog* (Figure S4I, J).

Taken together, these results demonstrate previously unappreciated heterogeneity within the satellite cell compartment and implicate Notch signaling within specific myogenic cell subpopulations.

### NOTCH2 signaling prevents myogenic differentiation *in vitro*

Given the existence of a specific satellite cell subpopulation enriched in NOTCH2 expression, we used primary satellite cells in a myotube-formation assay combined with Notch receptor antagonist antibodies to determine the contribution of NOTCH2 receptor for satellite cell proliferation and differentiation *in vitro*. When grown in culture, primary satellite cells enter the cell cycle and start dividing within 48 hours of culture. Within 5 days they start differentiating and fuse to form myotubes (Figure 3A, quantification in Figure 3B). By contrast, sustained stimulation of Notch signaling in satellite cells with bead-coupled recombinant Notch ligand inhibits differentiation and preserves a proliferative state. Blocking all Notch signaling, either with the gamma-secretase inhibitor DAPT or with a cocktail of antagonist anti-NOTCH1, 2 and 3 antibodies (αN1, αN2 and αN3) (Wu et al. 2010; Lafkas et al. 2015), abolished the effect of ligand-coated beads and resulted in myotube formation. When blocked individually, only inhibition of NOTCH2 resulted in myotube formation, showing that NOTCH2 is the principle receptor that acts to maintain satellite cells in a self-renewal and proliferative state *in vitro* and prevents differentiation. Blocking NOTCH1 or NOTCH3 individually had no effect. NOTCH2 inhibition also resulted in reduced expression of satellite cell marker genes *Pax7* and *Myf5* sustained by ligand beads (Figure 3C). Notably, expression of Notch target genes *Hey1, Hey2* and *Heyl* was increased when satellite cells were grown on bead-coupled recombinant Notch ligand, but decreased upon NOTCH2 inhibition, consistent with diminished Notch signaling. Taken together, these results indicate that NOTCH2 is the major Notch signaling transducer that mediates satellite cell maintenance *in vitro*.

**Figure 3.**
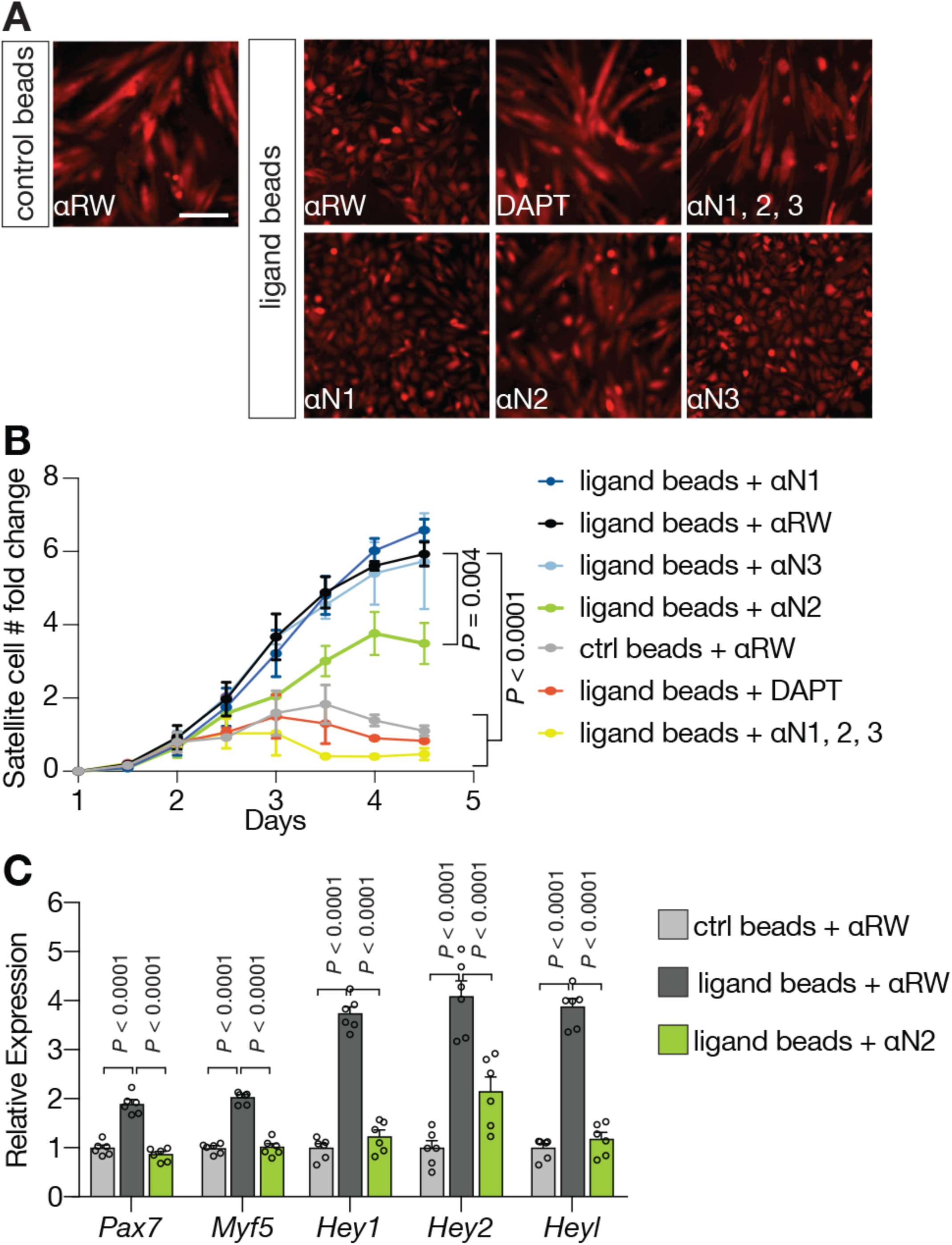
NOTCH2 signaling prevents myogenic differentiation *in vitro*. (A) Representative images of satellite cells after 4.5 days in culture grown either on control beads or ligand-coated beads, in the presence of control (RW) or anti-NOTCH antagonist antibodies as indicated. Scale bar, 100 μm. (B) Quantification of satellite cells numbers over 4.5 days (see Methods) (n = 6, biological replicates; *P* values two-way ANOVA at 4.5 days, compared to ligand beads + ⍺RW control, error bars, SEM) (C) Levels of satellite cell marker and Notch downstream target mRNAs were measured by qPCR normalized to *Rn18s*, under conditions as indicated. (*n* = 6, biological replicates, *P* two-way ANOVA with Dunnett’s multiple comparison correction, error bars, SEM).

### DLL1 and NOTCH2 are not required for satellite cell activation *in vivo*

To better understand roles of DLL1 and NOTCH2 during muscle regeneration and understand the roles of the subpopulations identified, we comprehensively investigated the function of the two Notch components *in vivo*. We performed a panel of experiments using antagonist antibodies targeting either DLL1 (*α*D1) or the NOTCH2 negative regulatory region (*α*N2) and compared their effect to isotype control antibody (*α*-Ragweed, *α*RW) at various time points during regeneration after cardiotoxin-mediated injury to the tibialis anterior (TA) muscle (Figure 4A and 5A). Systemic antibody treatment was initiated 1 day prior to muscle injury. Of note, anti-Notch antagonist antibodies used in these experiments are highly specialized therapeutic antibodies that had previously been developed and extensively characterized to specifically antagonize only one Notch receptor or ligand with similar high affinities, thus enabling the discrimination of individual functions (Wu et al. 2010; Lafkas et al. 2015; Ridgway et al. 2006; Tran et al. 2013). We first analyzed muscle at 2 days post-injury and found no difference in myogenic cell size or number (Figure 4B-C). Similarly, *Pax7*, *Myf5*, *Myod1* or *Myog* transcription factor expression in the muscle and total muscle weight were unchanged (Figure S5A). By contrast, downstream Notch signaling targets were downregulated (Figure S5A) indicating attenuated Notch signaling upon antibody treatment. These results indicate that DLL1- or NOTCH2-mediated signaling does not impair satellite cell activation after injury.

**Figure 4.**
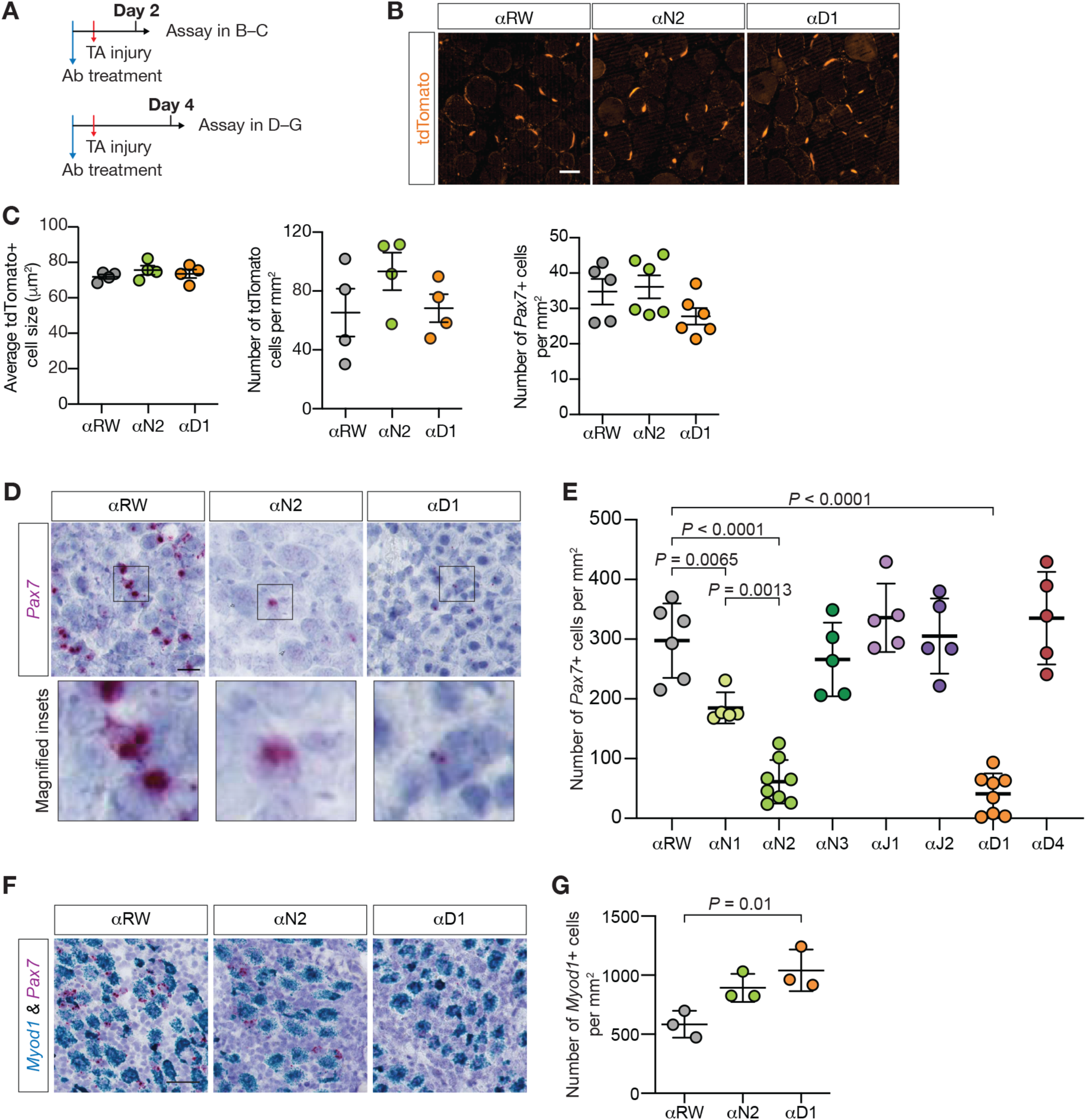
NOTCH2 and DLL1 signaling is required for satellite cell self-renewal *in vivo*. TA muscle in *Pax7.Cre_tdTomato* mice was cardiotoxin-injured and tissues were collected at 2 or 4 days post-injury. To inhibit DLL1 or NOTCH2-mediated signaling, specific antagonist antibodies were used and compared to control anti-Ragweed antibody. (A) Experimental schemes for *in vivo* muscle injury experiments with antagonist anti-Notch antibodies. (B) Representative images of cardiotoxin-injured TA sections at 2 days post-injury. Scale bar, 40 μm. (C) Cell size and number of tdTomato+ cells, and number of *Pax7+* cells does not change upon DLL1 or NOTCH2 inhibition at 2 days post-injury under indicated conditions. Each dot is average value from one mouse; error bars, SEM. (D) DLL1 and NOTCH2 inhibition decreases *Pax7* expression at 4 days post-injury. Representative images from ISH to detect mRNA expression of *Pax7* (pink) are shown. Magnified inset is shown to depict level of *Pax7* expression in cells. Scale bar, 20 μm. (E) DLL1 and NOTCH2 inhibition results in robust decrease in the number of *Pax7+* cells at 4 days post-injury. Number of *Pax7*+ cells was quantified under control or Notch blocking conditions (each dot is the average value from one mouse; *P* ordinary one-way ANOVA test, with Dunnett’s multiple comparisons test; error bars, SEM). (F) Representative images from multiplex ISH to detect mRNA expression of *Myod1* (blue) and *Pax7* (pink) at 4 days post-injury. Scale bar, 40 μm. Quantification (shown in G) shows an increase in *Myod1*+ cell numbers at day 4. Each dot represents average from one mouse. *P* ordinary one-way ANOVA test, with Dunnett’s multiple comparisons test, error bars, SEM.

**Figure 5.**
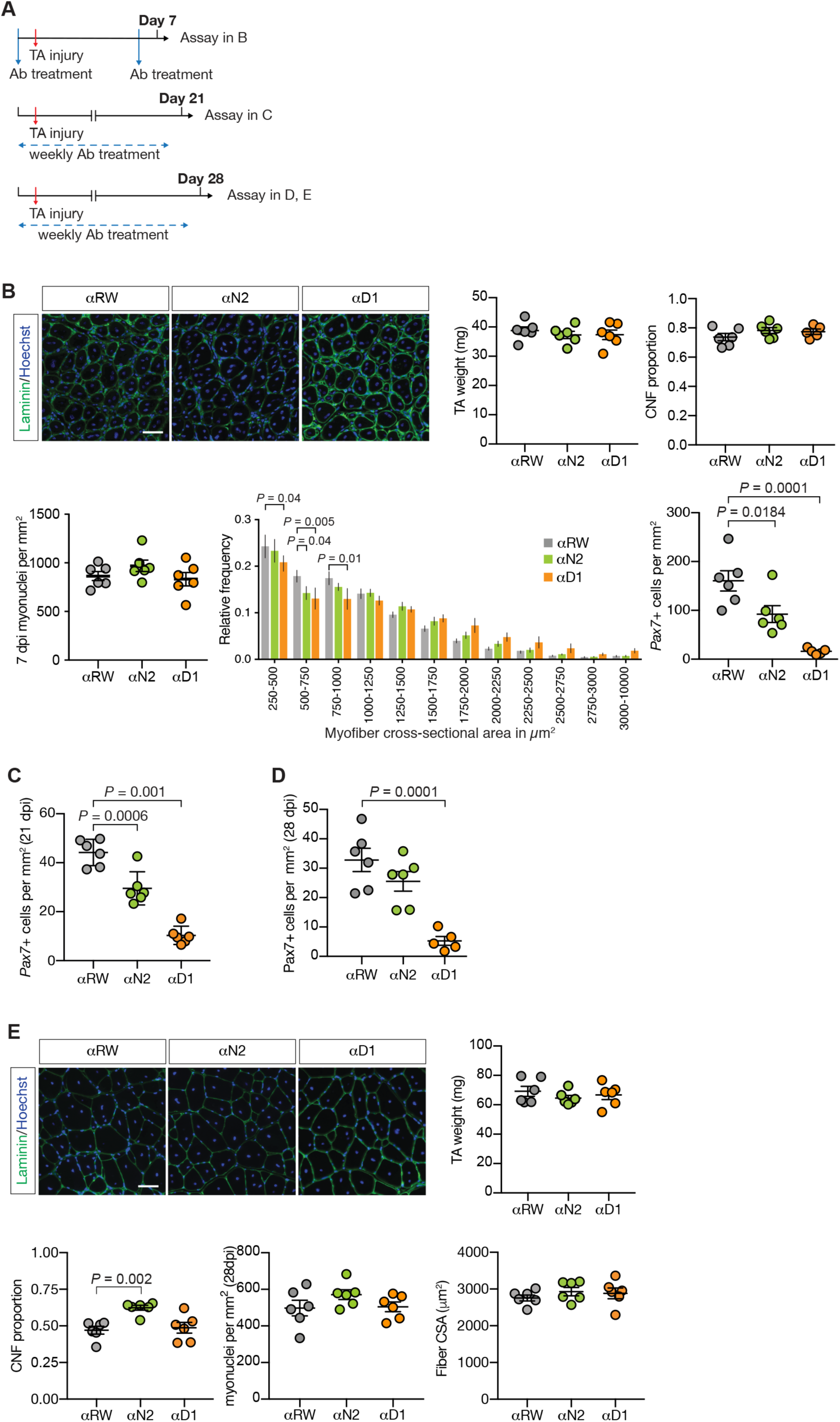
DLL1 and NOTCH2 signaling regulates satellite cell self-renewal. TA muscle in *Pax7.Cre_tdTomato* mice was cardiotoxin-injured and tissues were collected at 7, 21 or 28 days post-injury. To inhibit DLL1 or NOTCH2-mediated signaling, specific antagonist antibodies were used and compared to control anti-Ragweed antibody. (A) Experimental schemes for *in vivo* muscle injury experiments with antagonist anti-Notch antibodies. (B) Representative images of muscle sections at 7 days post-injury under indicated conditions. Laminin (green) was used to label muscle fibers and Hoechst stain (blue) was used to visualize nuclei. Total TA weight, the proportion of fibers with centrally localized nuclei (centrally nucleated fibers, CNF), number of myonuclei per area and relative frequency distribution of myofiber cross-sectional area was analyzed. Satellite cells were labeled with RNAscope probe for *Pax7* and their number was quantified and normalized to surface area. (*n* = 6 mice per condition, *P* two-way ANOVA test, with Dunnett’s multiple comparisons test, error bars, SEM). (C) At 21 days post-injury there is a significant decrease in the number of *Pax7+* cells. Satellite cells (stained with anti-*Pax7* RNAscope probe) were counted and normalized to section area at day 21 (*n* = 6 mice, *P* ordinary one-way ANOVA with Dunnett’s multiple comparison test, error bars, SEM). (D) Similar to (C) except at 28 days post-injury. (E) Representative images of muscle sections at 28 days post-injury under indicated conditions. Laminin (green) was used to label muscle fibers and Hoechst stain (blue) was used to visualize nuclei. Total TA weight, the proportion of fibers with centrally localized nuclei (centrally nucleated fibers, CNF), number of myonuclei per area and relative frequency distribution of myofiber cross-sectional area (CSA) was analyzed. (n = 6 mice per condition, *P* ordinary one-way ANOVA with Dunnett’s multiple comparisons test, error bars, SEM).

### DLL1 and NOTCH2 are required for sustained *Pax7* expression during regeneration

At 4 days post-injury, we observed that the number of *Pax7*+ cells was dramatically decreased upon DLL1 or NOTCH2 inhibition (Figure 4D, 4E, S5D). Not only was the overall number of positive cells significantly decreased, but the amount of *Pax7* RNA per cell appeared to be lower, as assessed by *in situ* hybridization signal (see insets in Figure 4D). Similarly, the number of *Myf5*+ cells was significantly decreased (Figure S5B). To investigate whether this effect is specific to DLL1 and NOTCH2, we utilized antagonist antibodies for other Notch receptors and ligands (NOTCH1, *α*N1; NOTCH3, *α*N3; JAG1, *α*J1; JAGGED2, *α*J2; and DELTA4, *α*D4) (Figure 4E, S5C, S5D). Treating mice with Notch antagonist antibodies and quantifying *Pax7* expression showed that inhibition of NOTCH1 also resulted in a decrease of *Pax7+* cell number, albeit more modestly than inhibition of NOTCH2 or DLL1, suggesting that there might be some redundancy between NOTCH2 and NOTCH1 in regulating *Pax7* expression *in vivo*. DLL1 was the only ligand demonstrating an effect on *Pax7*+ cell number. Concomitant with the decrease in the number of *Pax7*+ cells, inhibition of DLL1 resulted in a significant increase in the number of *Myod1+* cells at day 4 (Figure 4F, G). Of note, the number of caspase 3-positive cells was not increased, indicating that the decrease in *Pax7+* cells was not due to apoptosis (Figure S3E).

### DLL1 and NOTCH2 signaling regulates satellite cell self-renewal

To assess muscle differentiation and early muscle fiber formation under DLL1- and NOTCH2-blocking conditions, we analyzed muscles at 7 days post-injury (Figure 5B). The dramatic decrease in *Pax7*+ cell numbers at day 4 did not have any major effects on fiber formation as measured by the proportion of fibers with centrally localized nuclei, the number of myonuclei per area or the overall muscle weight at day 7. Measurement of the myofiber cross-sectional area pointed to in a possible shift towards larger myofibers. Similar to day 4, the number of *Pax7+* cells was significantly decreased. These observations prompted us to investigate whether satellite cells are able to self-renew during DLL1 and NOTCH2 inhibition. We analyzed their numbers at days 21 and 28 when the muscle regeneration cycle is nearly complete and observed a significant decrease at both time points (Figure 5C, D). Examination of fiber properties showed that there was no difference in fiber size or total muscle weight at day 28 (Figure 5E). NOTCH2 inhibition however, seemed to result in continued contribution of myogenic cells to myofibers, evidenced by a higher proportion of fibers with centrally localized nuclei at day 28. Overall, these results indicate that under the conditions of DLL1/NOTCH2 inhibition satellite cells are capable of successfully performing activation and differentiation processes, however self-renewal is impaired. When DLL1/NOTCH2 signaling is inhibited, satellite cells fail to maintain *Pax7* expression and progress towards myogenic differentiation, resulting in decreased self-renewal and re-establishment in the niche.

To further confirm these findings, we performed three sequential muscle injuries to TA muscles in mice treated with the antagonist antibodies (Figure 6, Figure S6). Inhibition of DLL1 dramatically impaired muscle regeneration upon triple cardiotoxin injury, as evident by a decrease in muscle weight, number of regenerating fibers (with centrally localized nuclei), smaller fiber size and increased muscle fibrosis (Collagen I deposition and macrophage infiltration) (Figure 6A, S6B). Inhibition of NOTCH2 had a more moderate effect on muscle regeneration, likely because of the weaker effect on satellite cell numbers. Importantly, inhibition of DLL1 resulted in a near-complete elimination of satellite cells (Figure 6B). Taken together these results indicate that satellite cell exhaustion in response to blocking of DLL1/NOTCH2 signaling results in failed muscle regeneration.

**Figure 6.**
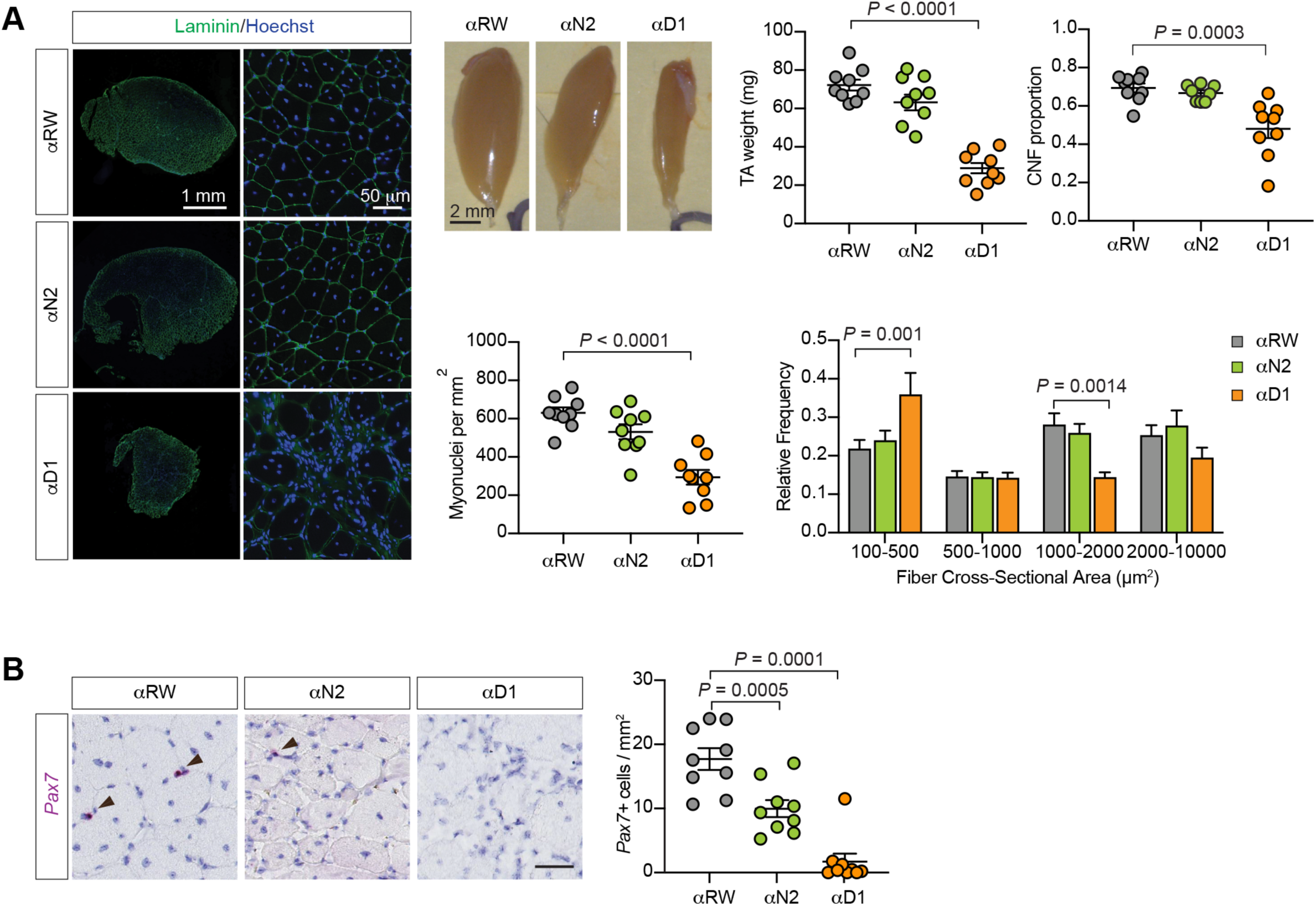
DLL1/NOTCH2 inhibition leads to impaired muscle regeneration upon multiple injuries. TA muscles were cardiotoxin-injured 3 consecutive times, every 25 days, and tissue was analyzed 25 days after the last injury. Throughout the experiment mice received weekly doses of antagonist *α*D1, *α*N2 or control *α*RW antibodies. (A) Under DLL1 inhibition conditions triple muscle injury leads to failure in regeneration, as evidenced by lower muscle weight, fewer regenerating muscles with centrally localized nuclei (centrally nucleated fibers, CNF), fewer myonuclei per section area and a shift towards smaller fibers. (n = 9 mice, *P* ordinary one-way ANOVA with Dunnett’s multiple comparisons test, error bars, SEM). (B) Under DLL1 and NOTCH2 inhibition muscles have fewer satellite cells upon multiple injuries. (n = 9 mice, *P* ordinary one-way ANOVA with Dunnett’s multiple comparisons test, error bars, SEM. Scale bar, 40 μm).

### *Notch2*-high cells balance myogenic differentiation and self-renewal

To understand satellite cell fate decisions controlled by DLL1/NOTCH2 signaling during injury, we performed single-cell RNA sequencing of satellite cells under DLL1- or NOTCH2-blocking or control (*α*RW) conditions at day 0 and day 4. Satellite cells at day 0 after *α*RW or *α*NOTCH2 treatment resembled those at day 0 without antibody treatment (Figure S7, Figure 2B). Myogenic cells at day 4 under control *α*RW condition showed a similar distribution across states as in the original data set (Figure 7A, 7B). By contrast, *α*DLL1 or *α*NOTCH2 treatment at day 4 resulted in a marked increase in proliferation and differentiation, at the expense of primary and ECM populations (Figure 7A, 7B). Biological replicate samples showed similar changes, with a stronger effect associated with DLL1 inhibition compared to inhibiting NOTCH2. The percentage of *Notch2*-high cells remained more stable across conditions, indicating that *Notch2*-high satellite cells do not depend on NOTCH2 signaling for maintenance.

**Figure 7.**
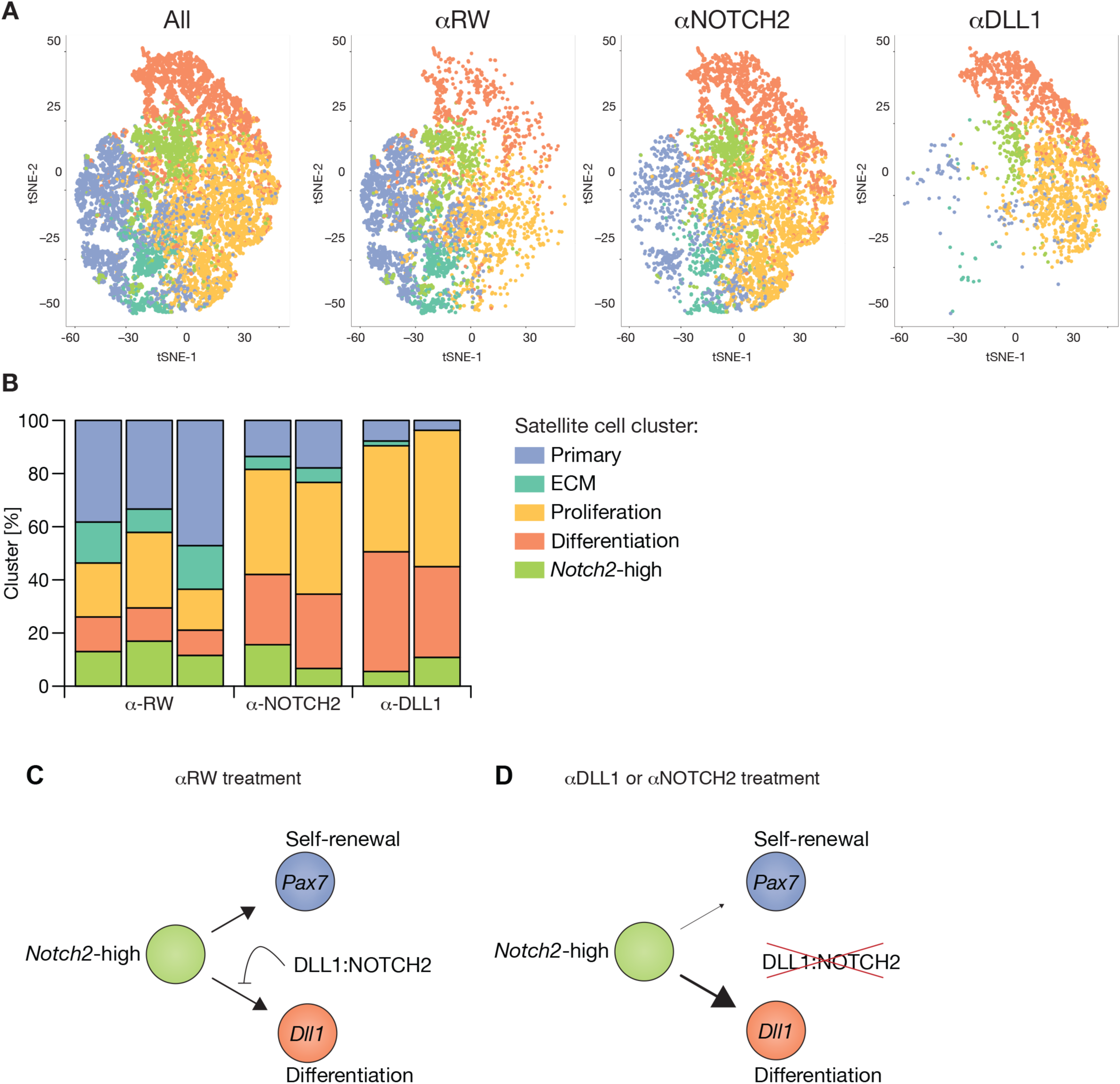
DLL1/NOTCH2 signaling controls satellite cell self-renewal. (A) t-SNE plot of single-cell expression profiles after antibody treatment at day 4 post-injury (n = 15,728). Cluster colors are detailed in the legend for panel (B). (B) Bar charts for the distribution of cells across satellite cell states at day 4 post-injury after treatment with *α*RW (n = 3), *α*NOTCH2 (n = 2), *α*DLL1 (n = 2). Cluster colors are same in (A) and (B). (C) Model of DLL1/NOTCH2 signaling controlling satellite cell self-renewal. G1-phase *Notch2*-high cells divide into cells with capacity to differentiate or self-renew. DLL1-expressing differentiating cells activate NOTCH2 signaling to inhibit myogenic differentiation and promote satellite cell self-renewal. (D) Disruption of DLL1/NOTCH2 signaling results in increased myogenic differentiation and lack of self-renewal.

To understand the effect of DLL1/NOTCH2 inhibition on individual myogenic populations, we performed differential gene expression analyses comparing *α*DLL1 or *α*NOTCH2 to *α*RW within each population (Figure S8A). We detected few expression changes within primary, proliferating and differentiating populations, suggesting the main consequence of DLL1/NOTCH2 inhibition is a shift in the distribution of cells across populations, rather than perturbing their expression profiles. By contrast, the *Notch2*-enriched population showed increased levels of differentiation markers and reduced levels of primary satellite cell markers (Figure S8A). Consistent with the observed gene expression changes, the previously seen heterogeneity within the *Notch2*-high cells (Figure S3A) underwent a shift from primary and ECM to differentiation and proliferation, mirroring the changes observed for the whole myogenic compartment (Figure S8B).

Taken together, these observations indicate that DLL1/NOTCH2 inhibition results in a shift in the cell fate of *Notch2*-high cells towards differentiation at the expense of self-renewal and ECM. Our results suggest a model by which DLL1 on differentiating cells stimulates NOTCH2 in a feedback loop to provide signaling required for triggering synthesis of ECM components and *Pax7* expression.

## DISCUSSION

### Satellite cell heterogeneity

Signals that govern satellite cell self-renewal have long remained elusive, despite intense research interest. A major obstacle for identifying such signals has been the inability to study this process *in vivo*. In this study we undertook a series of experiments to enable better understanding of satellite cell heterogeneity and mechanisms that regulate self-renewal in their native niche, the whole muscle tissue.

By employing single-cell RNA sequencing we detected heterogeneity among satellite cells during homeostasis. In particular, we identified a *Notch2*-enriched subpopulation, which persists during muscle regeneration. Heterogeneity of satellite cells during homeostasis has been discussed previously. However, segregation of isolated satellite cells was performed based on predetermined criteria (for example rate of division, level of Pax7, Pax3, Mx1, Myod1 expression) (Ono et al. 2012; Rocheteau et al. 2012; Der Vartanian et al. 2019; Scaramozza et al. 2019, Dell’Orso et al. 2019). Here we performed unbiased profiling of satellite cells during homeostasis and observed that they clustered within 2 subpopulations. The *Sdc4/*Notch2-enriched subpopulation reported in this study could also be detected in previously independently generated satellite cell single-cell RNA profiles (Giordani et al. 2019).

In addition to homeostasis, we report heterogeneity of satellite cells during the cell fate decision stage during regeneration. We describe 3 subpopulations of satellite cells, which provide insights into the mechanisms regulating self-renewal. Namely, we detect a transitional *Notch2*-high state which balances self-renewal and differentiation. We also detect an ECM-high and a primary state most closely related to quiescence. In addition to satellite cell heterogeneity, we describe a proliferating myoblast population and a *Dll1*-enriched differentiating population, revealing a switch from Notch receptor to Notch ligand expression upon commitment to differentiation. Expression data for each of these states provide a resource for further analysis of myogenic cell functions during regeneration.

An open question remains, whether all cell subpopulations present *in vivo* are equally amenable to the dissociation and purification procedure used in this study, in other words whether there exists an additional state which we were not able to purify. This question is pertinent not only to satellite cell profiling, but presents a challenge for single-cell studies in general, and future advances in cell isolation procedures will clarify any biases of the current state-of-the-art methods.

### Notch signaling in satellite cell self-renewal

The Notch signaling pathway plays important roles in muscle regeneration. Satellite cell-specific abolishment of Notch signaling leads to loss of satellite cells (Bjornson et al. 2012; Mourikis et al. 2012). Likewise, constitutive Notch signaling in cultured myogenic cells upregulates Pax7 and prevents differentiation (Wen et al. 2012). Further, a hypomorphic allele of *Delta like canonical Notch ligand 1* (*Dll1*) results in muscle hypotrophy resulting from precocious differentiation of progenitor cells during development (Schuster-Gossler et al. 2007). These studies undoubtedly indicated the importance of Notch signaling in satellite cells, but specific functions of individual Notch signaling components remained elusive. Given the vast regenerative potential of satellite cells, delineating specific signals that govern satellite cell self-renewal or differentiation *in vivo* is a crucial step towards harnessing this potential for medical purposes.

The role of specific Notch signaling components in muscle stem cell biology has been a major outstanding question in the field (Mourikis & Tajbakhsh 2014). In this work, we utilized Notch antagonist antibodies to delineate the function of specific receptors and ligands and identified the DLL1 ligand and NOTCH2 receptor as the principal mediators of satellite cell self-renewal in injured muscle. We show that inhibition of either DLL1 or NOTCH2 leads to a dramatic decrease in satellite cell marker expression and the number of satellite cells after injury. Our data demonstrate that DLL1 is the only Notch ligand necessary for satellite cell self-renewal. DLL1 inhibition leads to almost complete absence of satellite cells upon injury and failure to regenerate upon repeated injuries. Furthermore, Notch signaling has been reported to regulate Collagen V genes, extracellular matrix components, which in a cell-autonomous manner maintain quiescence of satellite cells (Baghdadi et al. 2018). Our data demonstrate that DLL1 inhibition leads to obliteration of the satellite cell population synthesizing ECM genes, positioning DLL1 as the furthest upstream self-renewal signal described to date.

Both NOTCH2 and NOTCH1 inhibition exhibited an effect on satellite cell self-renewal, though NOTCH1 effect was milder. This suggests a possible compensatory effect between Notch receptors in maintaining *Pax7* expression. We observed that satellite cells express NOTCH1 and NOTCH3 (Figure S2E). Interestingly, it was shown recently that satellite cells rely on both NOTCH1 and NOTCH2 to maintain a proliferative state and to prevent their differentiation when grown in culture, as well as to maintain quiescence *in vivo* (Fujimaki et al. 2018). Therefore, it is possible that NOTCH1 and NOTCH2 perform partially redundant functions. Notably, compared to control condition, NOTCH2 inhibition resulted in satellite cells contributing to muscle fibers at an increased rate during late regeneration at 28 days post-injury (Figure 5E). Together with the observation that *Notch2*-high cells are in G1 phase of the cell cycle, this suggests that NOTCH2 stimulation provides the required cellular environment for the *transition* into quiescence (thus quiescence establishment, not quiescence maintenance). We are only now starting to understand the differences between individual Notch components (Nandagopal et al. 2018) and future studies will further delineate signaling differences between stimulation of individual NOTCH receptors.

Our discovery of the ligand-receptor signaling pair that regulates satellite cell self-renewal also provides a better understanding of how proportional tissue regeneration is achieved. A previous report described that myogenic cells use remaining laminin, or “ghost fibers” to guide their migration (Webster et al. 2016). Our work further explains how this architecture is translated into a cell counting mechanism. When confined to the “ghost fiber” area, the DLL1-NOTCH2 axis provides a specific signal that triggers self-renewal, once sufficient cell density is achieved.

### Model for satellite cell self-renewal

Based on the data presented here, we propose a model whereby during muscle regeneration, the DLL1 ligand on differentiation-committed cells conveys a self-renewing signal to cells expressing the NOTCH2 receptor, with the balance between self-renewal and differentiation regulated by the presence or absence of neighboring DLL1-expressing cells (Figure 7C). DLL1 inhibition mimics a situation in which neighboring differentiating cells are absent, resulting in a shift towards more daughter cells committing to differentiation (Figure 7D). An outstanding question is whether the same feedback mechanism is employed during muscle growth. Given that *Dll1* is expressed in *Myog+* cells during the phase of postnatal muscle growth (Figure S4H) and that *Dll1* mutant embryos fail to develop muscle mass (Schuster-Gossler et al. 2007), it is plausible that the DLL1/NOTCH2 differentiation/self-renewal feedback loop is required for muscle growth as well.

It was recently demonstrated that in the skin epithelium, local differentiation drives epidermal stem cell self-renewal by a cell non-autonomous program (Mesa et al. 2018). These observations are strikingly similar to our findings in the muscle. While it was not shown which factors are involved in the skin epithelium, the physical interaction between differentiated cells and cells bound for self-renewal appears to be the hallmark of this mechanism. Our model depicts an elegant solution for stem cell self-renewal. Capitalizing on the spatial proximity between the differentiating myoblasts and satellite cells in the skeletal muscle compartment, the differentiating myoblasts use the highly conserved and dynamic Notch cell-to-cell signaling system to relay a self-renewal signal to satellite cells, and thus balance self-renewal and differentiation programs. Understanding the mechanisms that control stem cell self-renewal is a crucial step towards both correcting specific defects in disease, and harnessing the vast potential of adult stem cells for regenerative medicine.

## ACKNOWLEDGMENTS

We would like to thank the gRED Center for Advanced Light Microscopy (CALM) for imaging support, Jeffrey Eastham-Anderson for automating the SMASH software, the gRED FACS Core for cell sorting support, and Christopher R. Bjornson for scientific discussion. We thank Christopher Bohlen for providing feedback on the manuscript. We thank Casper Hoogenraad for his advice and input on the manuscript.

## AUTHOR CONTRIBUTIONS

A.J. conceptualized and supervised the project. V.Y. performed satellite cell isolations, FACS analyses, and *in vitro* experiments. M.C., L.K. and J.R. dosed animals and collected tissues for the *in vivo* experiments. J.R., V.Y. and A.J. processed tissues. A.J. and V.Y. analyzed *in vivo* experiments. O.F. performed automated analyses of *Pax7* areas. C.W.S. contributed the antibodies and discussed the results. Z.M. oversaw RNA-sequencing experiments and discussed the results. J.C. performed single-cell RNA sequencing library preparation. L.D.G. performed computational analysis of the data. A.J., L.D.G. and V.Y. formally analyzed the results. A.J wrote the original draft. A.J., L.D.G., V.Y. and Z.M. reviewed and edited the manuscript.

## METHODS

### *In vivo* studies

All animal studies were conducted in accordance with the Guide for the Care and Use of Laboratory Animals, published by the National Institutes of Health (NIH) (NIH Publication 8523, revised 1985). The Institutional Animal Care and Use Committee (IACUC) at Genentech reviewed and approved all animal protocols. All the mice were maintained in a pathogen-free animal facility under standard 12 h light/12 h dark cycle at 21°C with access to normal chow (Labdiet 5010) and water ad libitum. All the mice used for studies were 12-16 weeks old. All antibody doses administered were at a concentration of 20 mg/kg once seven days, except anti-Notch3 which was given at a dose of 30 mg/kg two times per seven days, and delivered via intraperitoneal injection.

Cardiotoxin injury was performed under isoflurane anesthesia. Animals were placed face up, injection site was prepped with alcohol, then injected with 50 μL of 10 μM Cardiotoxin (Cardiotoxin, *Naja pallida*, Millipore #217503) resuspended in sterile PBS. Needle was held parallel to the Tibia bone and introduced through the lower half of the muscle. By holding the needle parallel to the bone, inserted needle was guided towards the top half of the muscle. After placing the needle, cardiotoxin was slowly ejected while retracting the needle towards the entrance point. At the end of the injection needle was not pulled out immediately, but waited 2 – 3 seconds to prevent leakage of cardiotoxin. The relatively large volume of cardiotoxin injection ensured good tissue penetrance and injury throughout the TA. Cardiotoxin injuries consistently resulted in > 90% area damage when analyzed on sections made in the thickest middle part of the TA. Any part of tissue that was not damaged was excluded from the analysis.

### Tissue processing and analysis

TA muscles were isolated by first removing the skin covering the muscle. Skin was cut at the level of the proximal tendon and pulled up to completely expose the TA. Using forceps, fascia was carefully removed and TA was separated from the surrounding muscles. By cutting the tendons TA was isolated from the mouse. TA muscles were fixed in 2% PFA for 5 hours at room temperature and transferred into 30% sucrose solution at 4°C overnight. The next day muscles were placed on a cutting board and a straight cut was made at the distal end. TAs were place into Peel-A-Way Disposable Embedding Molds (S-22) (Polysciences, Inc. #18646A-1) in a vertical position and while supporting the top end with forceps embedder in Tissue-Tek O.C.T. Compound (VWR #4583). Vertically placed muscles were then frozen in isopentane cooled in liquid nitrogen. Blocks were cryosectioned to 8 μm perpendicularly to the vertical axes of the block, placed on Superfrost Plus Microscope Slides (Fisher #12-550-15), and kept at −80°C until use. To minimize possible effect of variation in angle of section, all muscles were isolated and frozen following same procedure, all histological analyses were done in a blinded manner and experiments were repeated. Per each animal minimum of 3 tissue sections were analyzed and average from all tissue sections from an animal is reported. Numbers of animals used per condition are indicated in figure legends.

*In situ* hybridization was performed using RNAscope^®^ 2.5HD Assay-RED (Advanced Cell Diagnostics #322360), RNAscope^®^ 2.5HD Assay-Duplex Assay (Advanced Cell Diagnostics #322430) or RNAscope^®^ Fluorescent Multiplex Assay v2 (Advance Cell Diagnostics # 323110) (Wang et al. 2012) according to the manufacturer’s protocols. Digital images of muscle sections stained by RNA Scope were acquired using the Nanozoomer 2.0-HT (Hamamatsu, Hamamatsu City, Shizuoka Pref, Japan) whole slide scanning system at x200 magnification. Quantification of *Pax7*+ or *Myod1+* cells was performed manually in blinded manner using Point Tool Counter in ImageJ (Schneider et al. 2012), normalized to surface of the injured area. Cells were counted only in the damaged area of muscle and any non-injured area was also excluded from surface measurement.

Regions of interest (ROI) outlining the muscle tissue were manually drawn and analyzed at full resolution using Matlab (Mathworks, Natick, MA). The images were converted into grayscale images and segmentation of tissue area was determined using intensity thresholding. The algorithm segmented out the tissue area present in the manually-drawn ROIs by thresholding for background in the grayscale-converted image.

To automatically quantify *Pax7* areas, within the ROIs *Pax7* areas were identified based on color thresholding for red, followed by minor morphological smoothing in the original RGB image.

To measure tdTomato-positive cell size and number at 2 days post-injury, muscle sections were analyzed using ImageJ (Schneider et al. 2012). Images were converted to 8-bit format, threshold set and “Analyze Particle” option used with the “size = 30-500 μm^2^” and “circularity = 0.00-0.90” with a custom macro script to automate. Cell counts reported by ImageJ were normalized to surface of injured area. Surface of injured area was measured using ImageJ, by first manually drawing around the area or injury, excluding any uninjured portion of the muscle.

RNAscope^®^ detects individual RNA molecules, which appear as distinct dots. To interpret *Notch2* staining pattern in non-injured muscle sections we used semi-quantitative scoring system based on number of dots per cell, as described in manufacturer’s protocol. High mRNA expression levels at 4 days post-injury precluded observation of individual dots in expressing cells and we did not perform quantification of individual RNA molecules in cells at 4 days post-injury.

To multiplex PAX7 immunohistochemistry with RNAscope, RNAscope protocol was first performed until detection step, followed by Avidin/Biotin Blocking Kit (Vector Labs SP-2001), Mouse on Mouse (M.O.M.) Blocking reagent (Vector Labs #MKB-2202), 1:5 anti-PAX7 antibody (DSHB, PAX7 supernatant) incubation overnight at 4°C, 1:200 dilution of anti-mouse biotin secondary (provided with Mouse on Mouse kit) for 20 minutes at room temperature (RT), 1:5000 dilution of Streptavidin-HRP (Abcam #ab7403) for 5 minutes at RT, 30 minute 0.3% Triton X-100 in PBS wash, and detected with Peroxidase Substrate Kit DAB (Vector Labs #SK-4100) for 20 minutes. Other antibodies used were anti-NOTCH2 (Cell Signaling, D67A6, 1:100 dilution), SPP1 (R&D Systems, AF808, dilution 1:1000), anti-RFP (Biorbyt, orb182397, dilution 1:250). Fluorescent secondary antibodies used were donkey anti-mouse Alexa Fluor 488 (Invitrogen, A-21202), donkey anti-rabbit Alexa Fluor 488 (Invitrogen, A-21206), donkey anti-goat Alexa Fluor 555 (Invitrogen, A-21432), all in dilution 1:400.

To measure muscle fiber size and centrally localized nuclei proportion, sections were stained with anti-Laminin antibody (DAKO #Z0097) and Hoechst, whole sections were imaged and 20X objective exports covering the entire section were quantified with SMASH (Smith & Barton 2014) with the following parameters: segmentation smoothing factor = 10, nuclear smoothing factor = 5, object smoothing factor = 10, minimum fiber area = 500 μm^2^, maximum fiber area = 10000 μm^2^, maximum eccentricity = 0.95, minimum convexity = 0.8, nuclear distance from border = 10 μm, minimum nuclear size = 20 μm^2^, and nuclear intensity threshold = 30.

Cleaved Caspase-3 staining required 5 minute treatment with Target Retrieval Solution (DAKO S1700), 1 hour room temperature incubation with Cleaved Caspase-3 (Asp175) Antibody (Cell Signaling #9661) at 1:200 dilution, 20 minute incubation with PolyVision Poly-HRP Anti-Rabbit IgG (Leica Biosystems #PV6119), and >1min detection with Peroxidase Substrate Kit DAB (Vector Labs #SK-4100). Anti-Type I Collagen-UNLB (SouthernBiotech #1310-01) and anti-F4/80 (Invitrogen #MF48000) staining were done at 1:40 and 1:100 concentrations respectively overnight in the cold and visualized with Alexa Fluor-conjugated secondary antibodies (Life Technologies) at 1:500 concentration.

### Satellite cell isolation

Satellite cells were isolated from uninjured hindlegs (day 0) or cardiotoxin-injured tibialis anterior muscles (day 4) of *PAX7.IRES.Cre.ki_Rosa26.LSL.tdTomato* (*PAX7.Cre_tdTomato*) mice. In this reporter strain satellite cells and all its progeny (myoblasts and muscle fibers) are labeled with tdTomato protein expression. Muscles cells were digested as in (Liu et al. 2015).

Satellite cells for single-cell RNA-seq profiling were purified with FACS for tdTomato/Calcein Blue-AM (ThermoFisher #C1429) double-positive cells. Based on results from our initial single-cell RNA-seq experiments, and results published during the course of our study (van den Brink et al. 2017), we recognized that this procedure can result in dissociation-related activation of stress response genes. In addition, we noticed numerous immune cells in the prep at 4 days post-injury (immune cells were not present in preps from non-injured muscles). To minimize these effects, we implemented the following improvements to our protocol: 1) 30 µM actinomycin D (Chemodex #A0043) was included during both enzyme digestion steps to prevent artifactual stress-response gene expression (Wu et al. 2017); 2) for single-cell RNA sequencing at 4 days post-injury, 320 µL Satellite cell isolation kit (Miltenyi Biotec #130-104-268) was used to deplete immune cells prior to FACS for every 4 TA muscles. These protocol improvements resulted in a higher number of captured satellite cells. Of 11 samples prepared for single-cell RNA sequencing, 4 were prepared without actinomycin D (untreated 0 dpi and 4 dpi samples presented in Figure 2, as well as one replicate each of the αRW and αN2 treated 4 dpi samples) (Figure S9A). We note that our observations pertaining to satellite cell heterogeneity were consistent across datasets, and were not affected by modifications to the cell isolation protocol. For myotube formation assay satellite cells were isolated using Satellite cell isolation kit from Miltenyi Biotec.

### Single-cell RNA sequencing

Sample processing for single-cell RNA sequencing was done using Chromium Single Cell 3’ Library and Gel bead kit v2 following manufacturer’s user guide (10x Genomics). The cell density and viability of single-cell suspension were determined by Vi-CELL XR cell counter (Beckman Coulter). The total cell density was used to impute the volume of single-cell suspension needed in the reverse transcription (RT) master mix, aiming to achieve ∼ 6,000 cells per sample. cDNAs and libraries were prepared following manufacturer’s user guide (10x Genomics). cDNA amplification and indexed libraries were prepared using 12 and 14 cycles of PCR, respectively. Libraries were profiled by Bioanalyzer High Sensitivity DNA kit (Agilent Technologies) and quantified using Kapa Library Quantification Kit (Kapa Biosystems). Each library was sequenced in one lane of HiSeq4000 (Illumina) to achieve ∼300 million reads following manufacturer’s sequencing specification (10x Genomics).

### Single-cell data processing

Single-cell RNA-seq data were processed with an in-house analysis pipeline. Briefly, reads were demultiplexed based on perfect matches to expected cell barcodes. Transcript reads were aligned to the mouse reference genome (GRCm38) augmented with the tdTomato reporter transgene sequence using GSNAP (2013-10-10) (Wu & Nacu 2010). Only uniquely mapping reads were considered for downstream analysis. Transcript counts for a given gene were based on the number of unique UMIs for reads overlapping exons in sense orientation. To account for sequencing or PCR errors, one mismatch was allowed when collapsing UMI sequences. Cell barcodes from empty droplets were filtered by requiring a minimum number of detected transcripts. Sample-specific cutoffs for cell detection were set to 0.1 times the total transcript count for the cell barcode at rank 30 (99th percentile for 3,000 cells). Data quality for individual libraries was assessed based on total read depth, percentage of reads with valid barcodes, percentage of demultiplexed reads in detected cells, number of detected cells, and number of analyzed cells (after removing non-myogenic cells as described in the following section) (Figure S9A). Sample quality was further assessed based on the distribution of per-cell statistics, such as total number of reads, percentage of reads mapping uniquely to the reference genome, percentage of mapped reads overlapping exons, number of detected transcripts (UMIs), number of detected genes, and percentage of mitochondrial transcripts (Figure S9B).

### Iterative cell filtering and clustering

For each sample, we filtered cells by iteratively clustering cells and removing clusters identified as non-myogenic cells or stressed cells based on the expression of known marker genes. Iterative cell filtering for two samples is illustrated in Figure S10. At most two filtering steps were performed for each sample. Cell clustering was performed using Seurat (2.0.0) (Macosko et al. 2015). A set of 41 genes was excluded to avoid identification of spurious clusters driven by genes with correlated expression due to overlapping exons. At each filtering step, the gene x cell matrix of raw UMI counts was log-normalized and scaled, regressing out total UMI count, and ‘S’ and ‘G2M’ cell cycle scores computed with Seurat function CellCycleScoring. Variable genes were identified with Seurat function FindVariableGenes. The top 10 principal components based on variable genes were used for clustering. We used clustering resolution 0.8 and 0.4 for iterative cell filtering and final clustering, respectively. For the Giordani dataset, iterative cell filtering resulted in selection of 769 cells used for analysis, final clustering was based on the top 3 principal components and resolution 0.4.

### Trajectory analysis

Pseudotime analysis was performed with Monocle2 (2.4.0) (Trapnell et al. 2014) using raw UMI counts for filtered cells as input, excluding cells with unusually low or high UMI count (outside two standard deviations from the mean on a log10 scale) and using genes with higher-than-expected dispersion and mean expression > 0.005 and < 5 for ordering cells.

### Identification of cell states

Clustering of filtered cells from the untreated 0 dpi and 4 dpi sample resulted in two and five clusters, respectively. Cluster marker genes were identified using Seurat function FindAllMarkers with default parameters. Pairwise cluster comparison was based on the overlap of identified markers genes and quantified by the Jaccard index (i.e. the number of unique markers identified for both clusters divided by the number of unique markers identified for either cluster). For the 4 dpi sample, the top 20 marker genes for each state were summarized into a summary score using Seurat function AddModuleScore. We used the R package glmnet (2.0-10) to fit a multinomial regression model using the five summary scores as input and cell states as output. The original untreated 4 dpi sample was used as training data, and cells in subsequent antibody-treated 4 dpi samples were assigned states and substates based on the most likely and second-most likely state, respectively.

### t-SNE of combined antibody-treated data sets

Filtered gene x cell matrices of raw UMI counts for antibody-treated 4 dpi samples were merged (*α*RW, n = 3; *α*N2, n = 2; *α*D1, n = 2). The merged matrix was log-normalized and scaled, regressing out total UMI count, and ‘S’ and ‘G2M’ cell cycle scores computed with Seurat function CellCycleScoring. Variable genes were identified with Seurat function FindVariableGene and principal components 1, 3, 4, 5 were used as input for t-SNE. Principal component 2 was associated with actinomycin D treatment and excluded from analysis.

### Differential expression analysis

We identified genes differentially expressed between *α*N2 (n = 2) and *α*RW (n = 3), and between *α*D1 (n = 2) and *α*RW (n = 3), separately for each state. ECM was excluded from analysis due to low prevalence of this state in *α*N2 and *α*D1 samples. For each state, gene expression profiles for individual cells from a particular sample were summarized by summing UMI counts across cells. This resulted in cell-state-specific pseudo-bulk expression profiles. Differential expression analysis was performed with limma-voom (Law et al. 2014). Briefly, we created a design matrix with four coefficients for the three antibody treatments as well as actinomycin D treatment. Genes with low expression were excluded by requiring a minimum CPM of 0.5 in at least two samples. Voom-transformed data were then analyzed with limma. We fit a linear model for all samples and subsequently extracted results for the two contrasts of interest. Moderated t-statistic p-values were adjusted for multiple testing using the Benjamini-Hochberg method.

### Myotube-formation assay

To generate Notch ligand-coupled beads, 500 μg BioMag® Goat anti-Human IgG (Fc Specific, Bangs Laboratories #BM562) were washed twice with DPBS without Calcium or Magnesium (Corning #21-031-CV) and incubated with 2 μg rat recombinant, Fc-fused JAG1 (R&D # 599-JD) overnight at 4°C with rotation. Beads were washed twice the next day and resuspended in 500 μL DPBS and used within 1 week. Bead-coupled-JAG1 ligand achieves non-selective activation of either NOTCH receptor expressed on cell surface, irrespective of the natural ligand (JAG1, 2 or DLL1, 3, 4) cells might find in *in vivo* setting. Satellite cells were plated on 96-well plate coated with ECM overnight at 4°C, incubated with 20 μg/mL antibodies for 24 hours before adding 10 μL ligand-bound bead suspension. Cells were imaged on the Incucyte Zoom v2016B using 20X objective for 5 days every 1 hour and tdTomato+ cell number quantified with Basic Analyzer Processing Definition to report the number of ‘red object count (1/well)’. Cell counts were normalized to the numbers measured at 24 hours when satellite cell numbers were stable following initial cell death.

### qPCR analysis

RNA was extracted from tibialis anterior muscles using RNAeasy Fibrous Tissue Kit (Qiagen #74704) and from satellite cells with Trizol reagent (ThermoFisher #15596026) according to manufacturer’s protocol. Reverse transcriptase reaction was performed using High-Capacity cDNA Reverse Transcription Kit (ThermoFisher # 4368814). cDNA was diluted 1:20 and qPCR reaction was performed with Taqman^TM^ Universal PCR Master Mix (ThermoFisher #4304437). Biological and technical triplicates were performed for each sample and relative expression with ΔΔCT method was measured on the ABI 7900HT instrument and analyzed with RQ Manager software v1.2.1 with *Rn18s* as reference control.

Taqman assays:

| Gene | Taqman Assay |
| --- | --- |
| *Rn18s* | Mm04277571_s1 |
| *Hes6* | Mm00517097_g1 |
| *Hey1* | Mm00516558_m1 |
| *Hey2* | Mm00469280_m1 |
| *Heyl* | Mm00516558_m1 |
| *Myf5* | Mm00435125_m1 |
| *Myod1* | Mm00440387_m1 |
| *Myog* | Mm00446194_m1 |
| *Pax7* | Mm01354484_m1 |

### Statistical analysis

PRISM (Graphpad Software (Prism Version 8)) was used to perform Student’s *t*-test and analysis of variance (ANOVA), as appropriate.

## FIGURE LEGENDS

**Figure S1.**
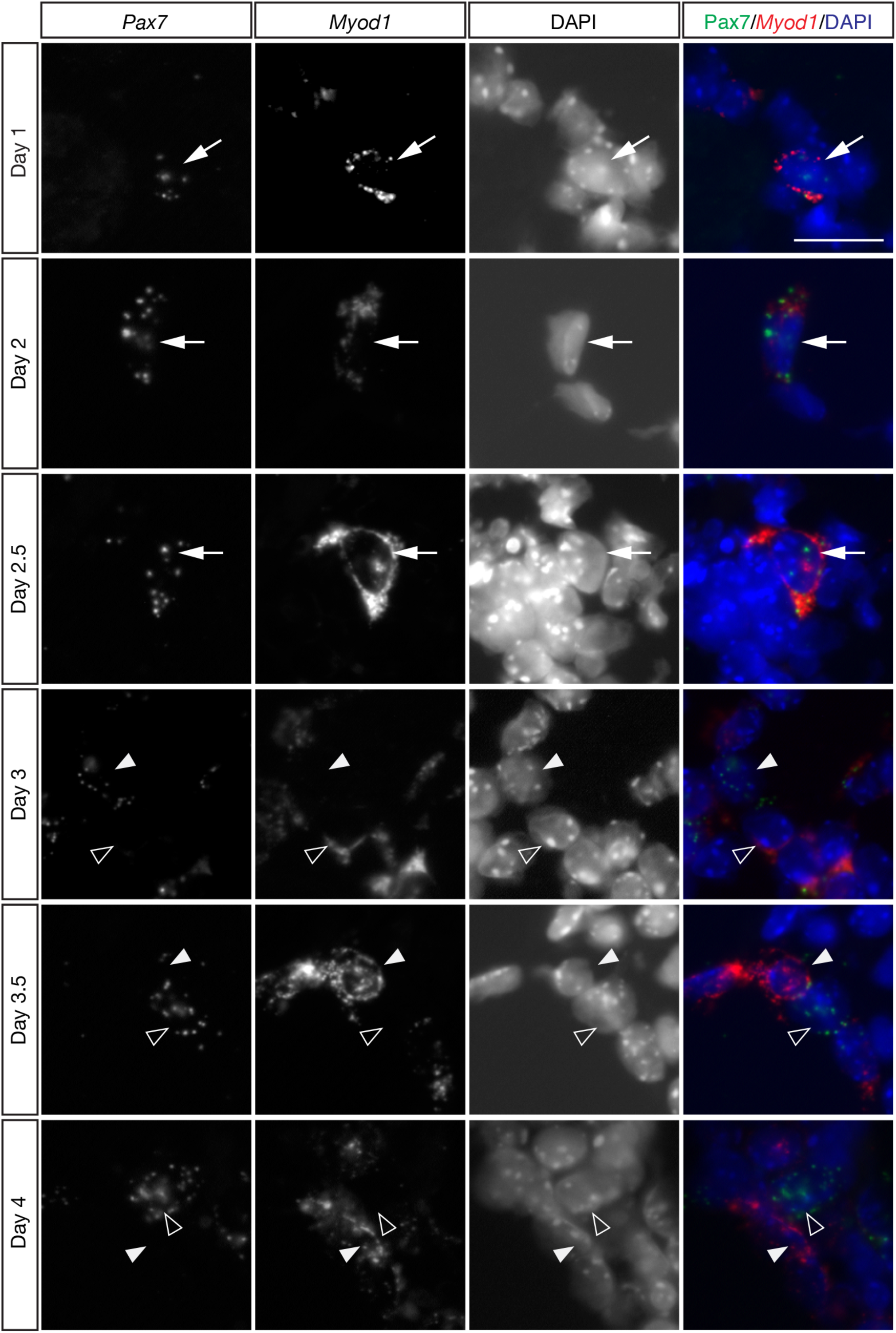
*In situ* analysis of myogenic transcription factor expression in early muscle regeneration. Related to Figure 1. Representative images of fluorescent *in situ* hybridization showing expression of myogenic factors *Pax7* and *Myod1* in cardiotoxin-injured tibialis anterior (TA) muscle at indicated time points post-cardiotoxin injury. At early time points (day 1 to 2.5) *Pax7* and *Myod1* are co-expressed in activated satellite cells. Starting at day 3 cells that are enriched in either *Pax7* or *Myod1* appear. At day 3.5 and 4 cells are either *Pax7*+ or *Myod1*+. Arrows point to double positive *Pax7+*/*Myod1+* cells, white arrowheads point to *Pax7*-enriched cells, empty arrowheads point to *Myod1*-enriched cells. Scale bar is 20 μm.

**Figure S2.**
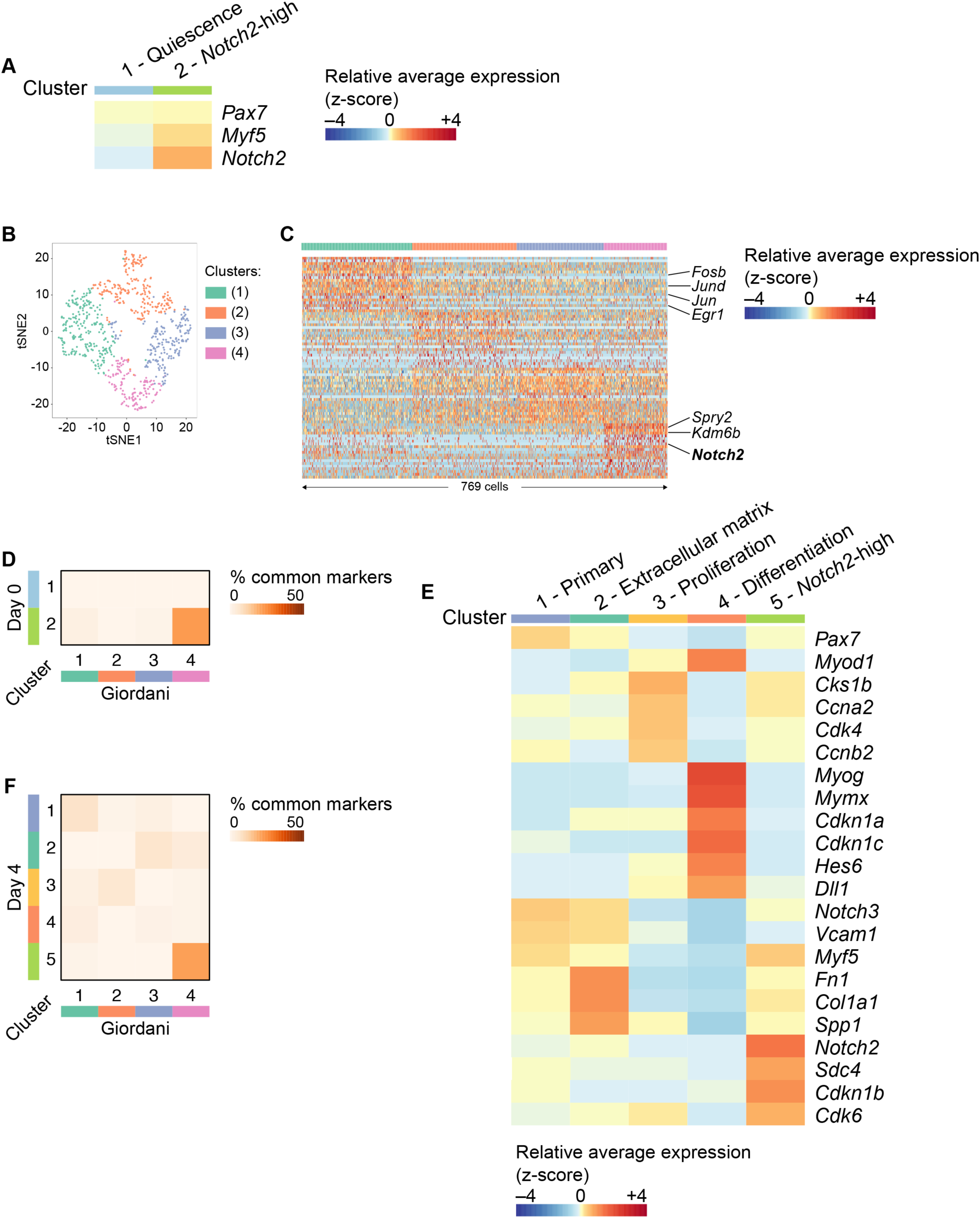
Expression of identified marker genes in satellite cell subpopulations. Related to Figure 2. (A) Heatmap showing average per-cluster expression of *Pax7*, *Myf5* and *Notch2* in satellite cell subpopulations during muscle homeostasis (no injury). (B) t-SNE plot of single-cell satellite cell expression profiles from Giordani et al. (2019). Cells clustered into four clusters, with cluster 4 showing overlapping markers with the *Notch2*-enriched cluster identified in our study. Cluster 1 showed enrichment in the expression of immediate-early genes indicative of a stress response, while clusters 2 and 3 showed weaker enrichment for individual genes and likely correspond to quiescent satellite cells with low transcriptional activity (our cluster 1). (C) Heatmap of top 20 identified marker genes for clusters shown in (B). Columns and rows correspond to cells and genes, respectively. Cells are ordered by cluster as shown in top sidebar, within-cluster order is random. (D) Cluster comparison between day 0 (our dataset) and Giordani et al. (2019) dataset. (E) Heatmap showing average per-cluster expression of selected genes at day 4. (F) Cluster comparison between day 4 (our dataset) and Giordani et al. (2019) dataset.

**Figure S3.**
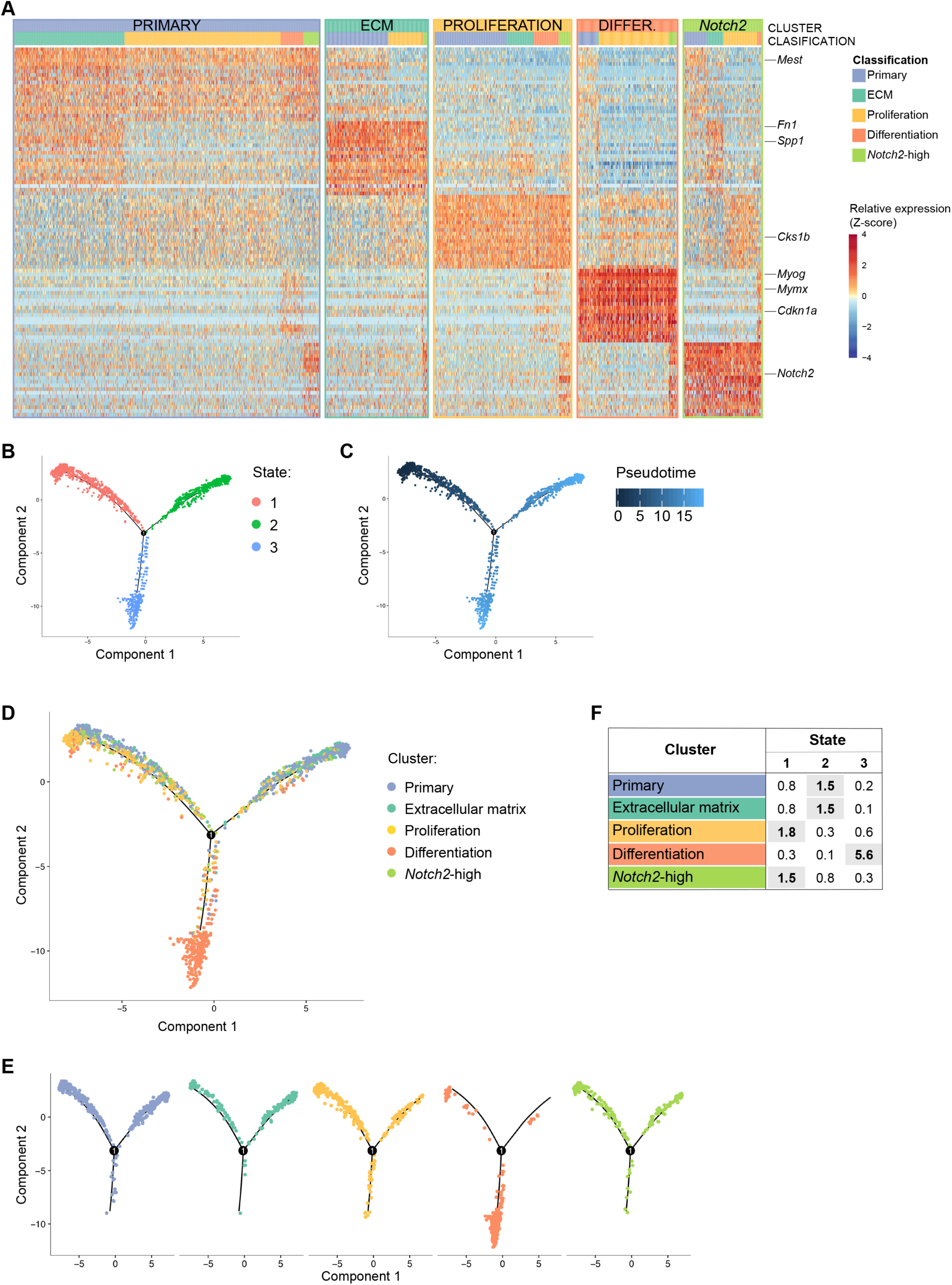
Satellite cell subpopulations represent dynamic states. Related to Figure 2. (A) Classification of cells into cellular states and substates as described in Methods. Heatmaps as in Figure 2D, with cells ordered by assigned state and substate. (B-F) Pseudotime analysis grouped cells into 3 different branches. Lines show the inferred trajectory, points show individual cells. (B) Cells colored by trajectory branch. (C) Cells colored by inferred pseudotime. (D, E) Cells colored by cluster shown as composite (D) or split by cluster (E). (F) Table with observed-over-expected cell count for each cluster and branch. Clusters enriched for a particular branch (> 1.0) are highlighted in bold.

**Figure S4.**
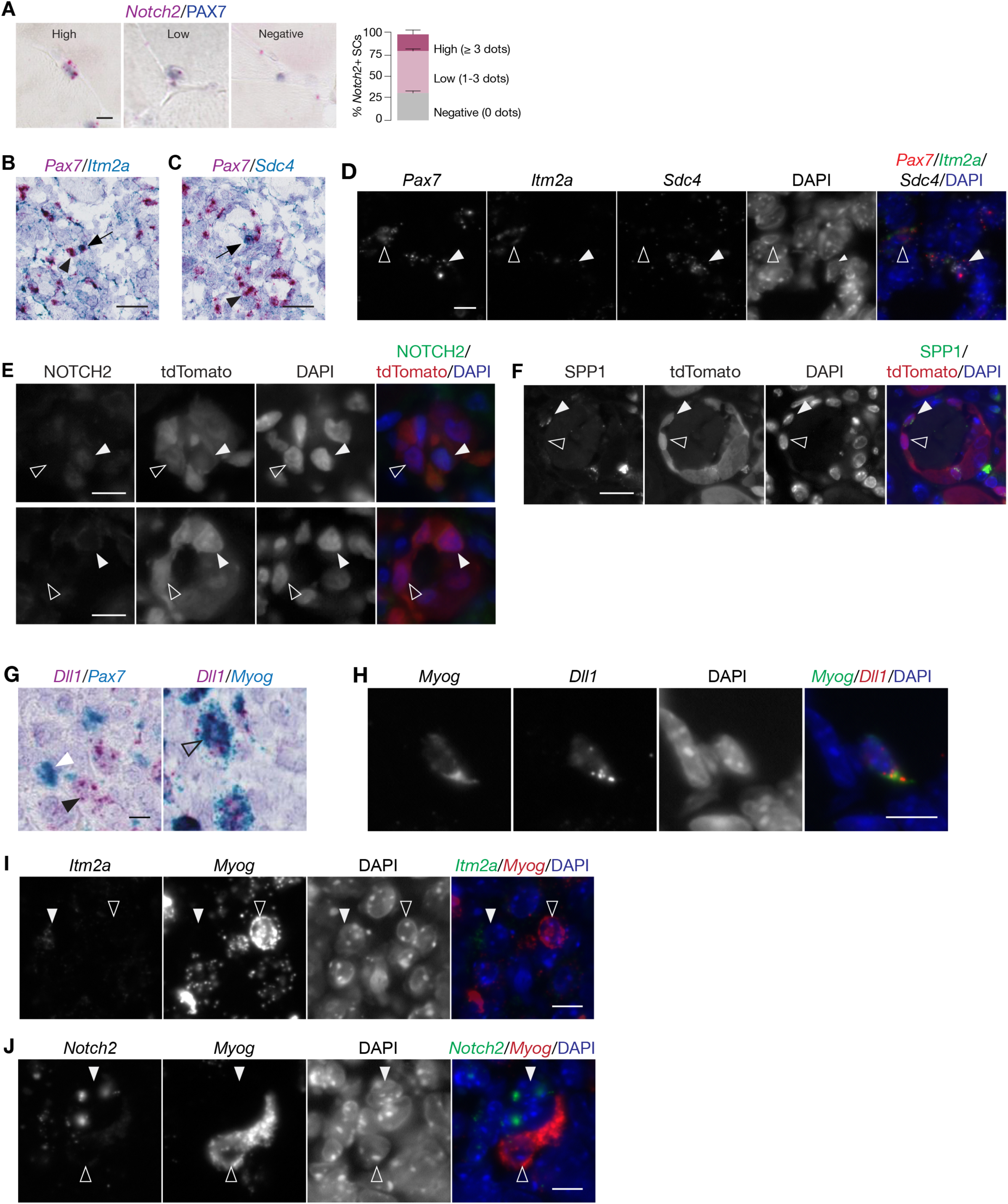
Characterization of cluster marker expression *in vivo*. Related to Figure 2. (A) Representative images of multiplexed RNAscope in situ hybridization (ISH) for *Notch2* (pink) and immunostaining for PAX7 (blue) in non-injured muscles. Quantification shows the percentage of PAX7+ cells per section that are *Notch2* “high” (3 or more dots), “low” (1-2 dots), and “negative” (0 dots), as described in Methods (n = 14 sections; error bars, SEM). Scale bar, 10 μm. (B-J) Representative images of multiplex *in situ* hybridization (ISH) or immunofluorescence staining of muscle tissue sections, all at 4 days post-injury (except H, which is non-injured TA from a 2-week-old mouse). (B) ISH staining using probes against *Pax7* (pink) and *Itm2a* (blue). Nuclei are stained with hematoxylin (purple). Arrow points to a satellite cells with enriched expression of *Itm2a*, arrowhead points to a cell with diminished *Itm2a.* Scale bar, 50 μm. (C) ISH staining using probes against *Pax7* (pink) or *Sdc4* (blue). Nuclei are stained with hematoxylin (purple). Arrow points to a satellite cells with enriched *Sdc4* expression, arrowhead points to a cell with diminished *Sdc4* expression. Scale bar, 50 μm. (D) ISH staining using probes against *Pax7* (red), *Itm2a* (green), *Sdc4* (white), and nuclei (DAPI, blue). Empty arrowhead points to a satellite cells with enriched *Itm2a* expression, white arrowhead to a satellite cell with enriched *Sdc4* expression. Scale bar, 10 μm. (E) Co-immunofluorescence staining of *Pax7.Cre_tdTomato* tissue sections using antibodies against NOTCH2 (green), tdTomato (red) and nuclei (DAPI, blue). White arrowheads indicate cells with NOTCH2 staining, empty arrowheads indicate cells negative for NOTCH2. Upper panel shows a cell with nuclear NOTCH2, lower panel shows a cell with cytoplasmic NOTCH2. Scale bar, 10 μm. (F) Co-immunofluorescence staining of *Pax7.Cre_tdTomato* tissue sections using antibodies against SPP1 (green), tdTomato (red) and nuclei (DAPI, blue). White arrowhead indicates cell with SPP1 staining, empty arrowhead indicates a cell negative for SPP1. Scale bar, 10 μm. (G) ISH staining using probes against *Dll1* (pink) and *Pax7* (blue) or *Myog* (blue), as indicated. White arrowhead points to a *Pax7+* cell, black arrowhead points to a *Dll1*+ cell, empty arrowhead points to a *Dll1*+/*Myog*+ cell. Scale bar, 10 μm. (H) ISH staining using probes against *Dll1* (red) and *Myog* (green) in non-inured muscle sections during postnatal growth phase (2-week-old mice). Scale bar, 10 μm. (I) ISH staining using probes against *Itm2a* (green) and *Myog* (red). Nuclei are stained with DAPI (blue). White arrowhead points to an *Itm2a+* cell, empty arrowhead points to a *Myog*+ cell. Scale bar, 10 μm. (J) ISH staining using probes against *Notch2* (green) and *Myog* (red). Nuclei are stained with DAPI (blue). White arrowhead points to a *Notch2+* cell, empty arrowhead points to a *Myog*+ cell. Scale bar, 10 μm.

**Figure S5.**
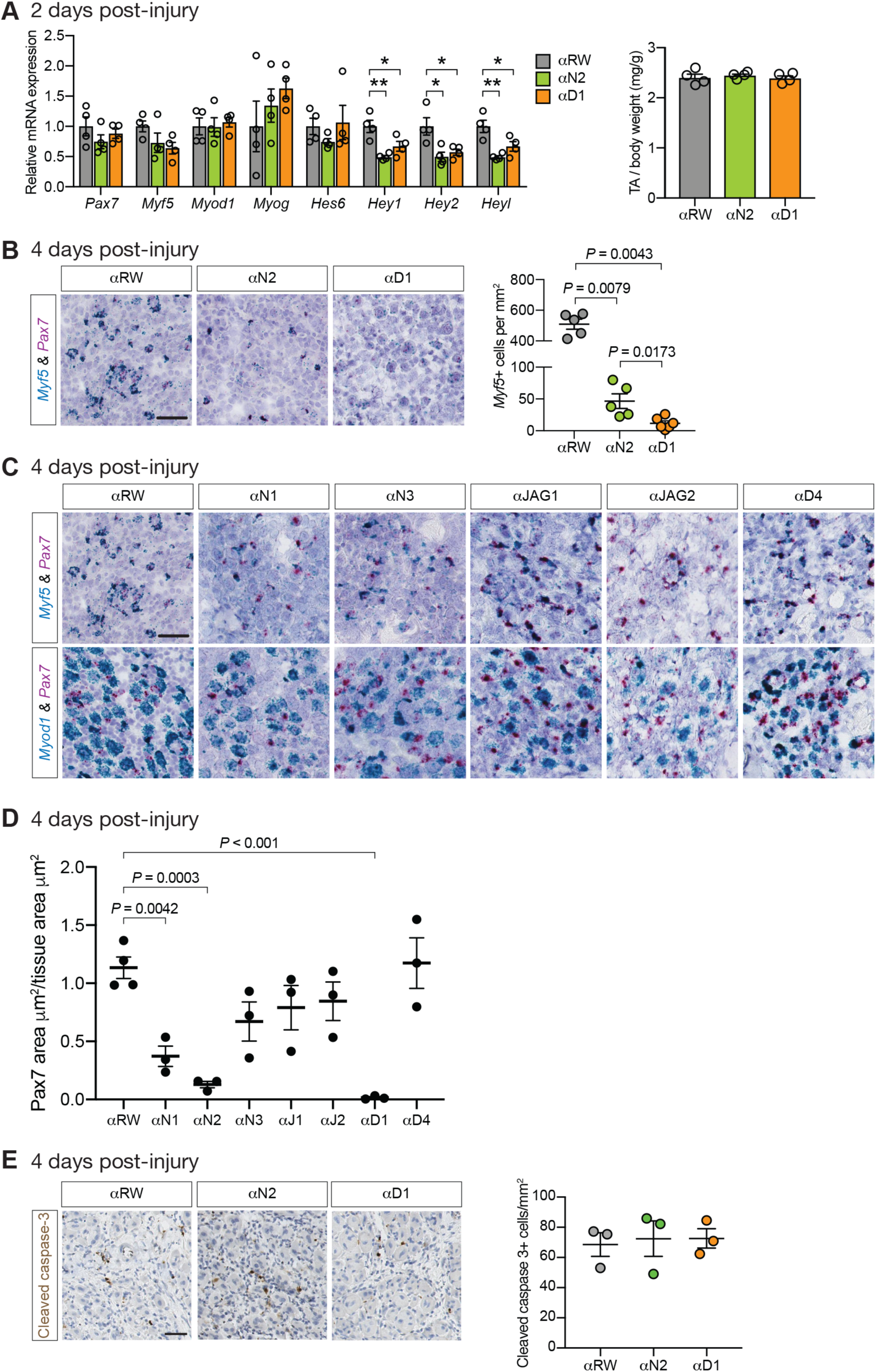
NOTCH2 and DLL1 does not affect satellite cell activation. Related to Figure 4. (A) TA of *Pax7.Cre_tdTomato* mice was injured with cardiotoxin and tissue was collected at day 2 post-injury. qPCR on TA muscle shows downregulation of Notch target genes under NOTCH2 or DLL1-inhibitory conditions when compared to control Ragweed (RW) (*n* = 3 mice per condition, *P* ordinary one-way ANOVA with Dunnett’s multiple comparison test, error bars, SEM). TA weight was unchanged (n = 4 mice per condition). (B) At 4 days post-injury there is a significant decrease in the number of *Myf5+* cells under NOTCH2- or DLL1-blocking conditions. Scale bar is 40 μm. *n* = 5 mice per condition. (C) Representative images of sections stained with RNAscope probes for *Pax7* (pink) and *Myf5* (blue) or *Myod1* (blue), under indicated conditions are shown. Scale bar is 40 μm. (D) Automated quantification of the size of all areas with red (*Pax7*-positive) signal within the total tissue area (*P* ordinary one-way ANOVA test, with Dunnett’s multiple comparisons test, error bars, SEM). (E) Number of caspase 3-positive cells was evaluated to test for increased cell death during muscle regeneration under NOTCH2- or DLL1-inhibitory conditions. Representative images are shown, no significant difference was observed between conditions, *n* = 3 mice per condition; error bars, SEM.

**Figure S6.**
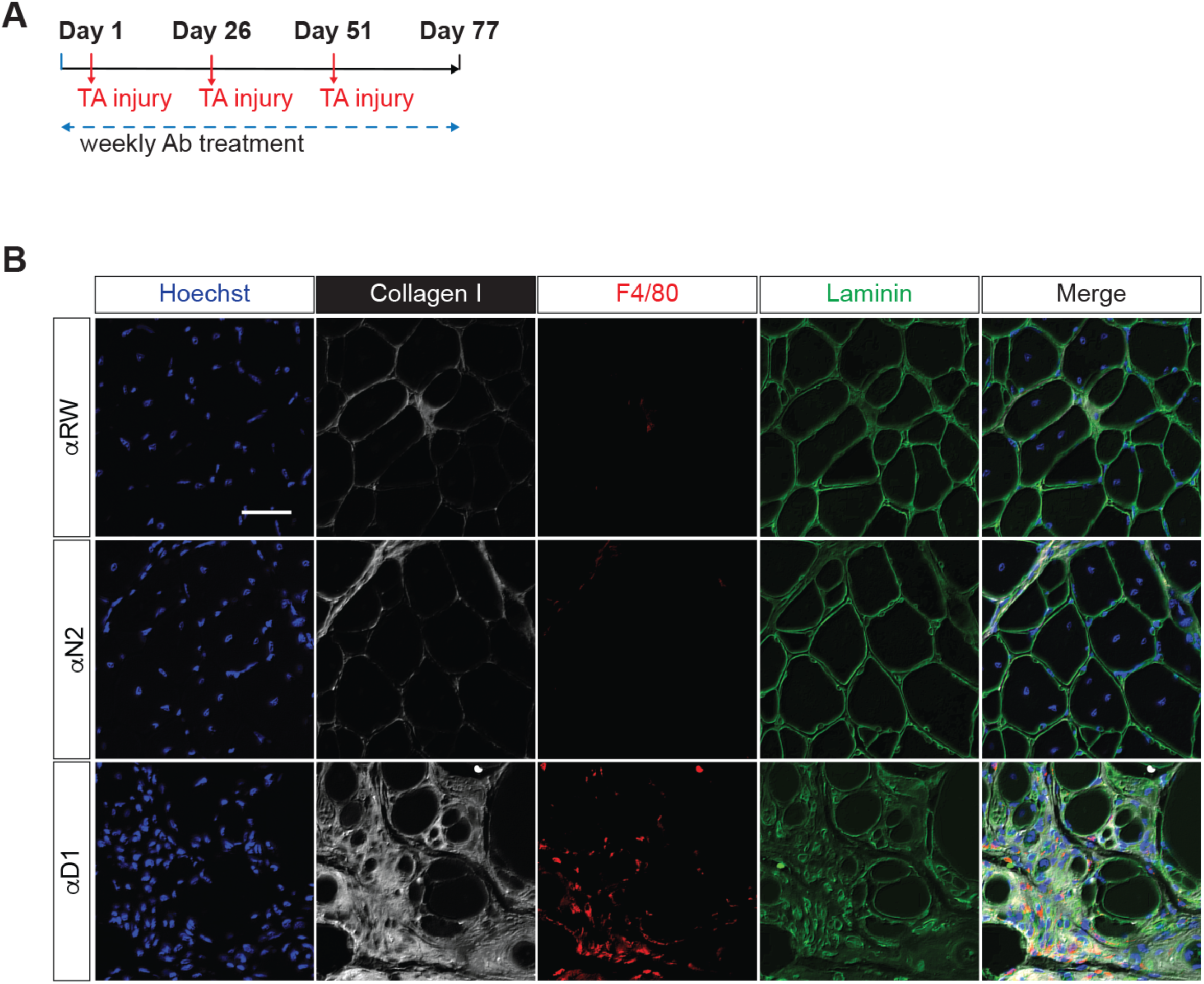
DLL1 inhibition leads to muscle fibrosis after 3 injuries. Related to Figure 6. (A) Experimental scheme for triple muscle injury experiment with antagonist anti-Notch antibodies. (B) Representative images of TA sections of cardiotoxin-injured TAs. labeled with Hoechst, Collagen Type I, the macrophage marker F4/80, and Laminin. Scale bar, 40 *µ*m.

**Figure S7.**
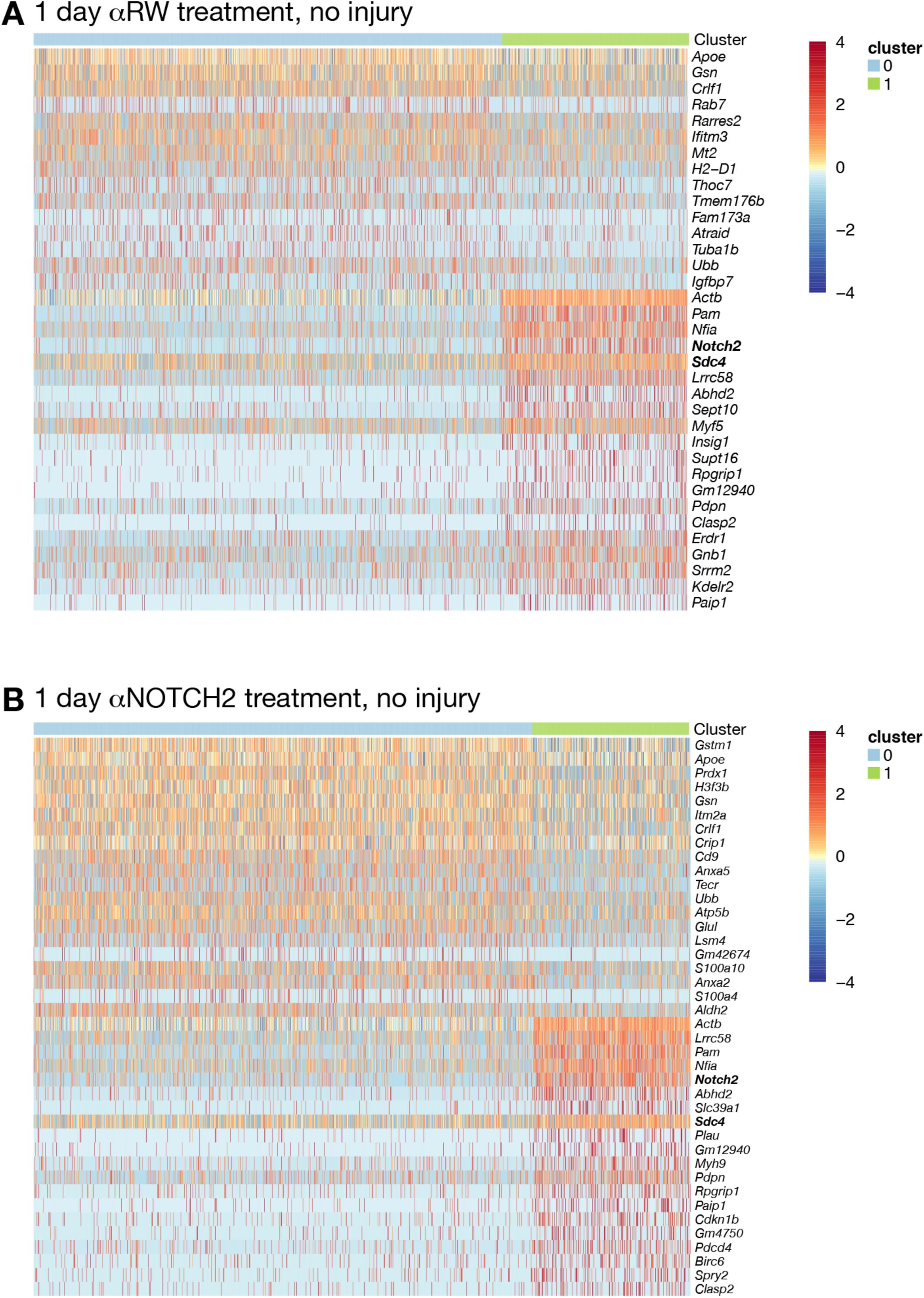
Satellite cell heterogeneity in non-injured TA. Related to Figure 7. Heatmaps of top 20 identified marker genes for clusters at day 0 under *α*RW (n = 1,679) (A) or *α*NOTCH2 condition (n = 1,447) (B). For cluster 1 under *α*RW condition (A) only 15 markers genes were identified. Otherwise as in Fig. 2B.

**Figure S8.**
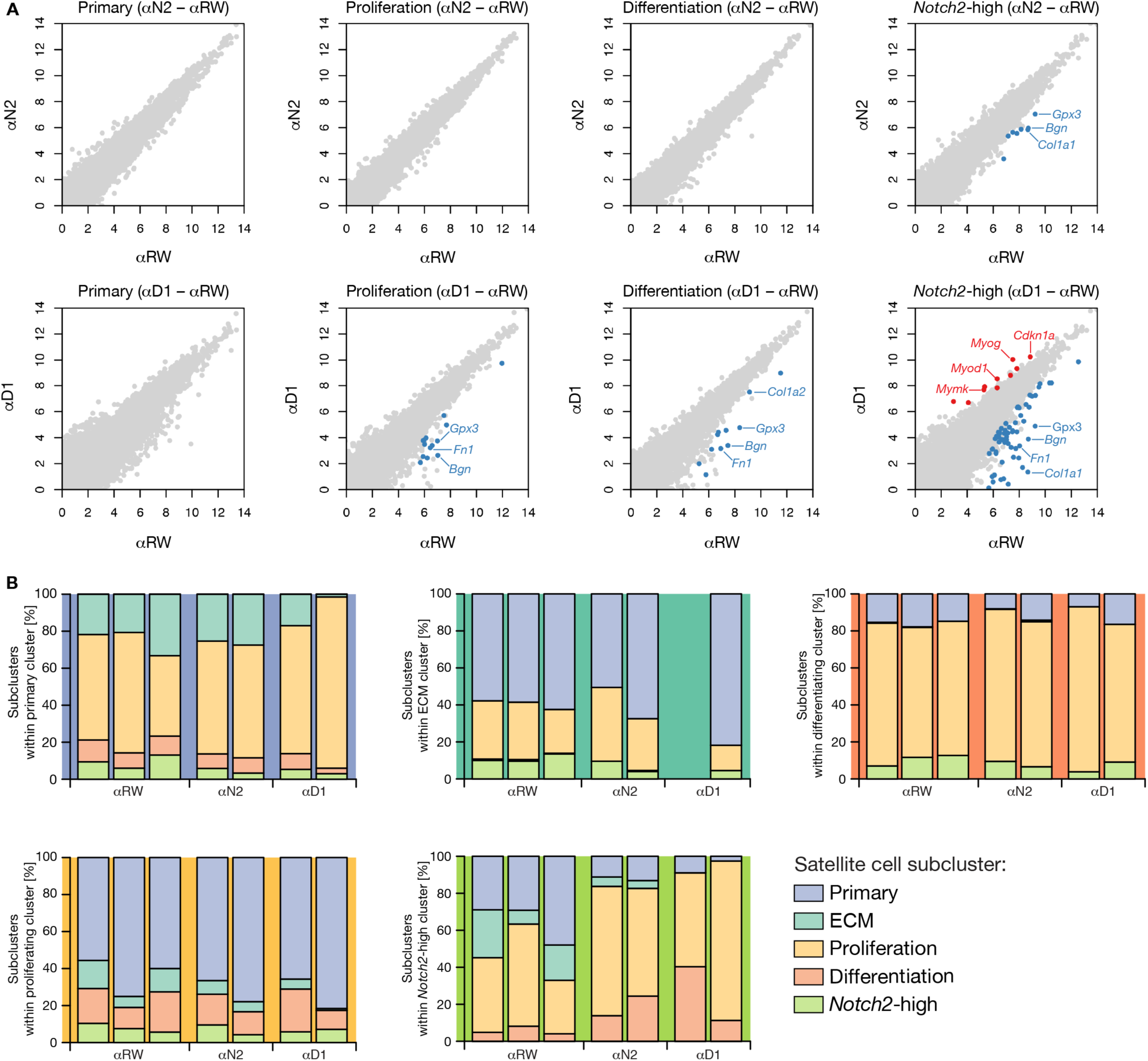
Gene expression changes following antibody treatment. Related to Figure 7. (A) Differential expression analysis for individual cell populations reveals gene expression changes within the *Notch2*-high subpopulation after *α*D1 antibody treatment. Genes with increased and decreased expression in *α*N2 or *α*D1 relative to *α*RW are highlighted in red and blue, respectively (adjusted p < 0.001, two-sided moderated t-test). Downregulation of ECM marker genes and upregulation of differentiation genes was observed. (B) Bar charts illustrating heterogeneity within satellite cell populations after treatment with *α*RW (n = 3), *α*N2 (n = 2) or *α*D1 (n = 2) antibodies. Separate panels show heterogeneity within primary, ECM, differentiation, proliferation and *Notch2*-high populations. One replicate for αD1 did not include cells classified as ECM.

**Figure S9.**
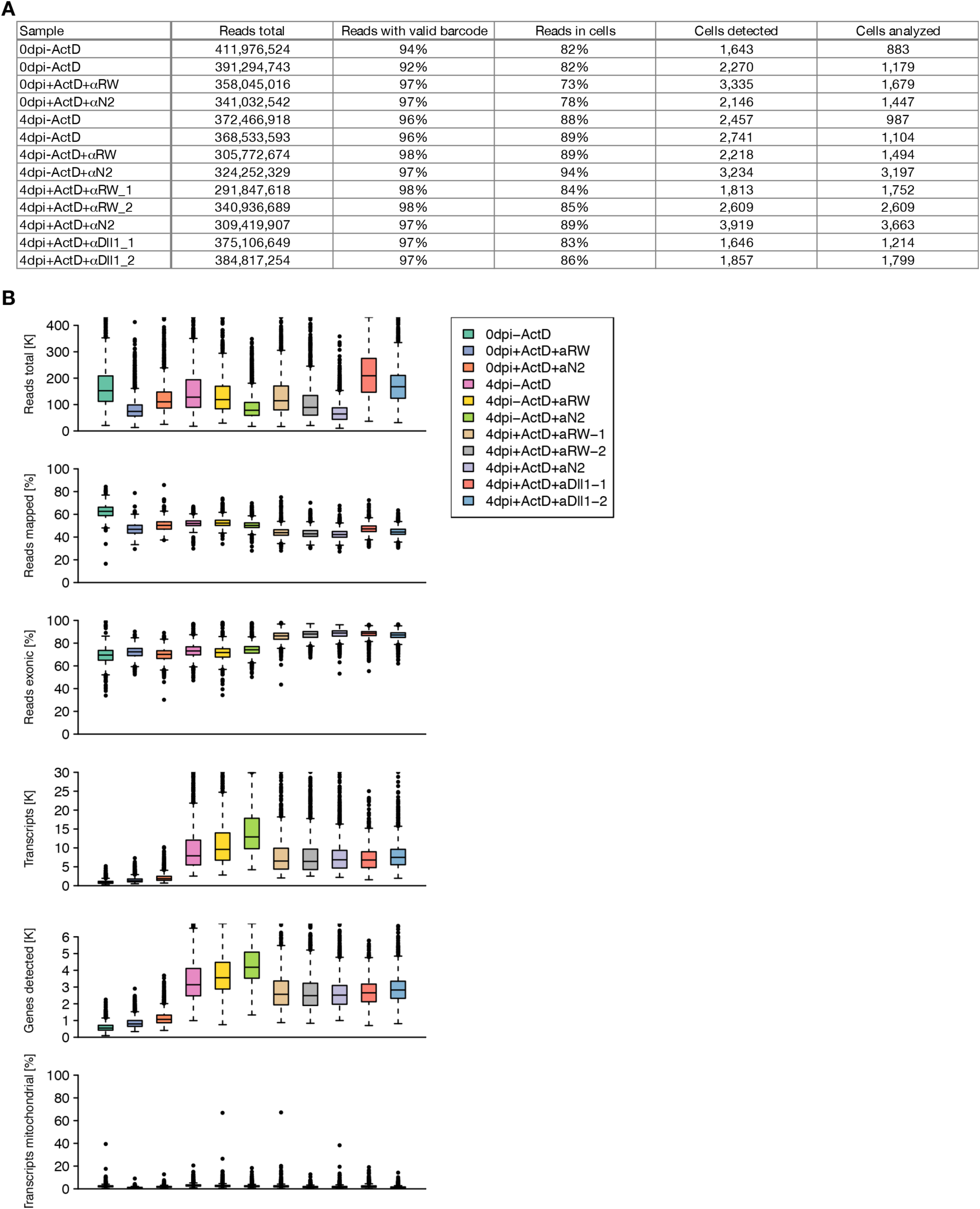
Single-cell data quality assessment. (A) Summary statistics for single-cell sequencing libraries used in this study, including total read depth, percentage of reads with valid barcodes, percentage of demultiplexed reads in detected cells, number of detected cells, and number of analyzed cells after removing non-myogenic cells. Two single-cell libraries were prepared for samples 0dpi-ActD and 4dpi-ActD. (B) Boxplots of per-cell statistics for analyzed cells for each sample. Panels show total reads, percentage of reads mapping uniquely to the reference genome, percentage of mapped reads overlapping exons, number of detected transcripts (UMIs), number of detected genes, and percentage of mitochondrial transcripts. Boxes indicate the interquartile range (IQR), center lines the median, whiskers extend to the most extreme data point within 1.5 × IQR from the box.

**Figure S10.**
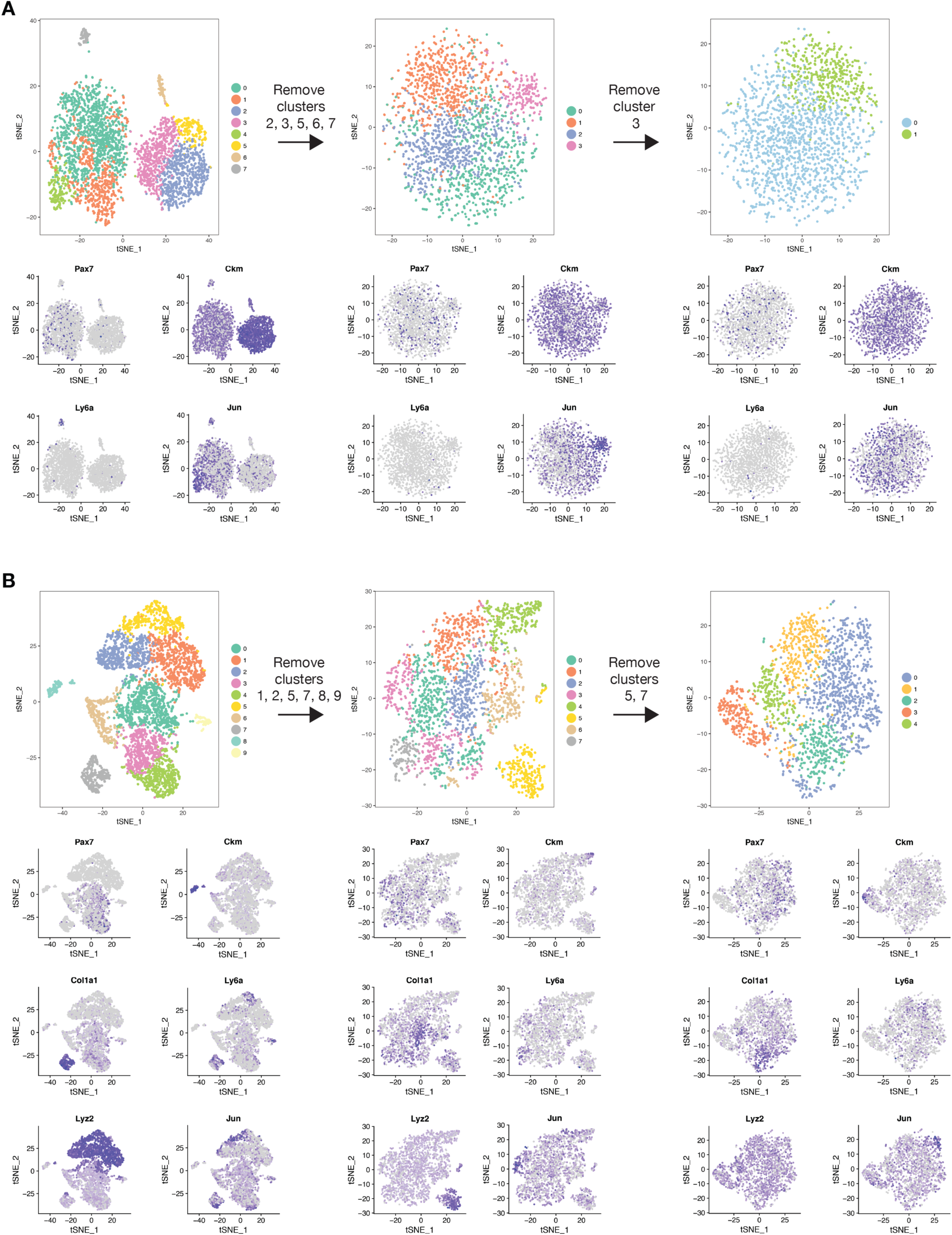
Removal of non-myogenic cells by iterative cell filtering. (A) Iterative cell filtering for untreated 0 dpi sample. After initial clustering, cells in clusters 2, 3, 5, 6 (muscle gene expression) and 7 (endothelial gene expression) were removed. After re-clustering, cells in cluster 3 (stress response gene expression) were removed. Final clustering resulted in two clusters. (B) Iterative cell filtering for untreated 4 dpi sample. After initial clustering, cells in clusters 1, 2, 5 (immune gene expression), 7 (tenocyte gene expression), 8 (muscle gene expression) and 9 (endothelial gene expression) were removed. After re-clustering, cells in cluster 5 (immune gene expression) and 7 (FAP gene expression) were removed. Final clustering resulted in five clusters.

## Supplementary Tables

**Table S1**. Identified marker genes for each cluster at day 0 post-cardiotoxin injury.

**Table S2**. Identified marker genes for each cluster at day 4 post-cardiotoxin injury.

**Table S3**. Differentially expressed genes within individual cell subpopulations after *α*-DLL1 or *α*-NOTCH2 treatment.

## References

Baghdadi, M.B. et al., 2018. Reciprocal signalling by Notch–Collagen V–CALCR retains muscle stem cells in their niche. Nature, 557(7707), pp.714–718.

Beauchamp, J.R. et al., 2000. Expression of CD34 and Myf5 defines the majority of quiescent adult skeletal muscle satellite cells. Journal of Cell Biology, 151(6), pp.1221–1233.

Bentzinger, C.F. et al., 2013. Fibronectin regulates Wnt7a signaling and satellite cell expansion. Cell Stem Cell, 12(1), pp.75–87.

Bi, P. et al., 2018. Fusogenic micropeptide Myomixer is essential for satellite cell fusion and muscle regeneration. Proceedings of the National Academy of Sciences, 115(15), pp.3864–3869.

Bjornson, C.R.R. et al., 2012. Notch signaling is necessary to maintain quiescence in adult muscle stem cells. Stem Cells, 30(2), pp.232–242.

Braun, T. & Gautel, M., 2011. Transcriptional mechanisms regulating skeletal muscle differentiation, growth and homeostasis. Nature Reviews Molecular Cell Biology, 12(6), pp.349–361.

Collins, C.A. et al., 2005. Stem Cell Function, Self-Renewal, and Behavioral Heterogeneity of Cells from the Adult Muscle Satellite Cell Niche. Cell, 122(2), pp.289–301.

Cornelison, D.D.W. & Wold, B.J., 1997. Single-cell analysis of regulatory gene expression in quiescent and activated mouse skeletal muscle satellite cells. Developmental Biology, 191(2), pp.270–283.

Cossins, J. et al., 2002. Hes6 regulates myogenic differentiation. *Development (Cambridge*, England), 129(9), pp.2195–207.

Fujimaki, S. et al., 2018. Notch1 and Notch2 Coordinately Regulate Stem Cell Function in the Quiescent and Activated States of Muscle Satellite Cells. Stem Cells, 36(2), pp.278–285.

Gao, X. et al., 2001. HES6 acts as a transcriptional repressor in myoblasts and can induce the myogenic differentiation program. Journal of Cell Biology, 154(6), pp.1161–1171.

Giordani, L. et al., 2019. High-Dimensional Single-Cell Cartography Reveals Novel Skeletal Muscle-Resident Cell Populations. Molecular Cell, 74(3), p.609–621.e6.

Giordani, L., Parisi, A. & Le Grand, F., 2018. Satellite Cell Self-Renewal 1st ed., Elsevier Inc.

Hasty, P. et al., 1993. Muscle deficiency and neonatal death in mice with a targeted mutation in the myogenin gene. Nature, 534(6437), pp.501–506.

Hawke, T.J. & Garry, D.J., 2001. Myogenic satellite cells: physiology to molecular biology. Am J Physiol Cell Physiol, 280, pp.1358–1366.

Hiramuki, Y. et al., 2015. Mest but not MIR-335 affects skeletal muscle growth and regeneration. PLoS ONE, 10(6), pp.1–15.

Kuang, S. et al., 2007. Asymmetric Self-Renewal and Commitment of Satellite Stem Cells in Muscle. Cell, 129(5), pp.999–1010.

Lafkas, D. et al., 2015. Therapeutic antibodies reveal Notch control of transdifferentiation in the adult lung. Nature, 528(7580), pp.127–131.

Lagha, M. et al., 2013. Itm2a Is a Pax3 Target Gene, Expressed at Sites of Skeletal Muscle Formation In Vivo. PLoS ONE, 8(5), pp.1–10.

Law, C.W. et al., 2014. voom: Precision weights unlock linear model analysis tools for RNA-seq read counts. Genome Biology, 15(2):R29.

Liu, L. et al., 2015. Isolation of skeletal muscle stem cells by fluorescence-activated cell sorting. Nature Protocols, 10(10), pp.1612–1624.

Macosko, E.Z. et al., 2015. Highly parallel genome-wide expression profiling of individual cells using nanoliter droplets. Cell, 161(5), pp.1202–1214.

Mauro, A., 1961. Satellite Cell of Skeletal Muscle Fibers. The Journal of Biophysical and Biochemical Cytology, 9(2), pp.493–495.

Mesa, K.R. et al., 2018. Homeostatic Epidermal Stem Cell Self-Renewal Is Driven by Local Differentiation. Cell stem cell, 23(5), pp.677–686.

Mourikis, P. et al., 2012. A critical requirement for notch signaling in maintenance of the quiescent skeletal muscle stem cell state. Stem Cells, 30(2), pp.243–252.

Mourikis, P. & Tajbakhsh, S., 2014. Distinct contextual roles for Notch signalling in skeletal muscle stem cells. BMC Developmental Biology, 14(1), pp.1–8.

Nandagopal, N. et al., 2018. Dynamic Ligand Discrimination in the Notch Signaling Pathway. Cell, 172(4), p.869–880.e19.

Ono, Y. et al., 2012. Slow-dividing satellite cells retain long-term self-renewal ability in adult muscle. Journal of Cell Science, 125(5), pp.1309–1317.

Ridgway, J. et al., 2006. Inhibition of Dll4 signalling inhibits tumour growth by deregulating angiogenesis. Nature, 444(7122), pp.1083–1087.

Rocheteau, P. et al., 2012. A subpopulation of adult skeletal muscle stem cells retains all template DNA strands after cell division. Cell, 148(1–2), pp.112–125. Available at: http://dx.doi.org/10.1016/j.cell.2011.11.049.

Scaramozza, A. et al., 2019. Lineage Tracing Reveals a Subset of Reserve Muscle Stem Cells Capable of Clonal Expansion under Stress. Cell Stem Cell, 24(6), p.944–957.e5.

Schneider, C.A., Rasband, W.S. & Eliceiri, K.W., 2012. NIH Image to ImageJ: 25 years of image analysis. Nature Methods, 9(7), pp.671–675.

Schuster-Gossler, K., Cordes, R. & Gossler, A., 2007. Premature myogenic differentiation and depletion of progenitor cells cause severe muscle hypotrophy in Delta1 mutants. Proceedings of the National Academy of Sciences, 104(2), pp.537–542. Available at: https://www.ncbi.nlm.nih.gov/pmc/articles/PMC1766420/pdf/zpq537.pdf.

Seale, P. et al., 2000. Pax7 Is Required for the Specification of Myogenic Satellite Cells. Cell, 102(6), pp.777–786.

Smith, L.R. & Barton, E.R., 2014. SMASH - semi-automatic muscle analysis using segmentation of histology: A MATLAB application. Skeletal Muscle, 4(21).

Tonami, K. et al., 2013. Calpain-6 Deficiency Promotes Skeletal Muscle Development and Regeneration. PLoS Genetics, 9(8), pp.1–11.

Tran, I.T. et al., 2013. Blockade of individual Notch ligands and receptors controls graft-versus-host disease. Journal of Clinical Investigation, 123(4), pp.1590–1604.

Trapnell, C. et al., 2014. The dynamics and regulators of cell fate decisions are revealed by pseudotemporal ordering of single cells. Nature Biotechnology, 32(4), pp.381–386.

Der Vartanian, A., et al., 2019. PAX3 Confers Functional Heterogeneity in Skeletal Muscle Stem Cell Responses to Environmental Stress. Cell Stem Cell, 24(6), pp.958–973.

Wang, F. et al., 2012. RNAscope: A novel in situ RNA analysis platform for formalin-fixed, paraffin-embedded tissues. Journal of Molecular Diagnostics, 14(1), pp.22–29.

Webster, M.T. et al., 2016. Intravital Imaging Reveals Ghost Fibers as Architectural Units Guiding Myogenic Progenitors during Regeneration. Cell Stem Cell, 18(2), pp.243–252.

Wen, Y. et al., 2012. Constitutive Notch Activation Upregulates Pax7 and Promotes the Self-Renewal of Skeletal Muscle Satellite Cells. Molecular and Cellular Biology, 32(12), pp.2300–2311.

Wu, T.D. & Nacu, S., 2010. Fast and SNP-tolerant detection of complex variants and splicing in short reads. Bioinformatics, 26(7), pp.873–881.

Wu, Y. et al., 2010. Therapeutic antibody targeting of individual Notch receptors. Nature, 464(7291), pp.1052–1057.

Wu, Y.E. et al., 2017. Detecting Activated Cell Populations Using Single-Cell RNA-Seq. Neuron, 96(2), pp.313–329. Available at: https://doi.org/10.1016/j.neuron.2017.09.026.

Zammit, P.S., 2008. All muscle satellite cells are equal, but are some more equal than others? Journal of Cell Science, 121(18), pp.2975–2982.

Zammit, P.S. et al., 2004. Muscle satellite cells adopt divergent fates: A mechanism for self-renewal? Journal of Cell Biology, 166(3), pp.347–357.

Zammit, P.S. et al., 2006. Pax7 and myogenic progression in skeletal muscle satellite cells. Journal of Cell Science, 119(9), pp.1824–1832.

